# Enhanced 4Pi single-molecule localization microscopy with coherent pupil based localization and light sheet illumination

**DOI:** 10.1101/586404

**Authors:** Sheng Liu, Fang Huang

## Abstract

Over the last decades, super-resolution techniques have revolutionized the field of fluorescence microscopy. Among them, interferometric or 4Pi microscopy methods exhibit supreme resolving power in the axial dimension. Combining with single-molecule detection/localization and adaptive optics, iPALM/4PiSMS/W-4PiSMSN allowed 10-15 nm isotropic 3D resolution throughout the whole cell. However, further improving the achieved 3D resolution poses significantly challenges which, in part, is blocked by the complexity of single-molecule emission pattern generated by these systems rendering a large portion of information carrying photons unusable. Here we introduce a localization algorithm that achieves the theoretical information limit for 4Pi based single-molecule switching nanoscopy (4Pi-SMSN), and demonstrates improvements in resolution, accuracy as well as applicability comparing with the state of art 4Pi-SMSN methods. Further, with a novel 4Pi-compatible light-sheet illumination reducing the fluorescence background by >5-fold, we demonstrated the new system enables further improvement in the achievable resolution of 4Pi/interferometric single-molecule imaging systems.

## Introduction

The achievable resolution of a far-field fluorescence microscope was constrained by the diffraction limit of light, approximately 200-300 nm in the lateral direction and 500-700 nm in the axial direction. Over the last decades, significant efforts were made to overcome this resolution limit. Based on confocal and widefield microscopy geometries respectively, 4Pi (type A-C) (*1-3*) and I^n^M (*4-6*) methods use coherent illumination and/or detection based on two opposing objectives to improve the resolution by 3-7 fold in the axial dimension (*7*). Combining coherent detection and the stochastic switching of single molecules with high precision localization, 4Pi (also known as interferometric) based single-molecule switching nanoscopy (SMSN) techniques, such as iPALM(*8*) and 4Pi-SMS (*9*), allow another 10 fold improvement in the axial resolution (*8, 10*) in comparison to conventional 3D SMSN. Further incorporating adaptive optics and interferometry-specific algorithm design, W-4PiSMSN (*11*) allowed high-resolution reconstruction of the whole mammalian cell (up to ∼9 μm) without deterioration of resolution throughout the imaging depth.

The resolution enhancing capacity of the interferometric systems comes from their complex point spread function (PSF) patterns, thereafter, referred as 4Pi-PSF. One of its distinct features is the rapid intensity modulation within the pattern center along the depth (axial) direction which, in turn, improves the axial resolution when combined with confocal, stimulated emission depletion microscopy and single-molecule localization microscopy methods. Aside from the center peak, the complexities of 4Pi-PSF majorly arise from the features such as rings and lobes at its periphery. Depending on the axial position of the single emitter, these features contains up to 80% of the entire 4Pi-PSF energy (**Materials and Methods**). Theoretical precision calculations based on Fisher information theory suggested that better localization precision can be obtained in both axial and lateral direction (*10, 12*) when compared with other incoherent dual objective systems. To date, these complex features remain largely unexplored when localizing single-molecule emission patterns from 4Pi systems, due to the challenge in modeling this complex pattern with high accuracy. An accurate 4Pi-PSF model must be able to take into account of the static imperfections of the interferometric single-molecule imaging system, such as aberrations and transmission variances in both interferometric arms, partial coherence due to the broadband emission spectra and the dynamic changes of the system, such as the temperature-dependent changes of the interferometric cavity length. Furthermore, due to the specific optical geometry of 4Pi systems (dual objective detection with large NA), difficulties in adapting existing light-sheet illumination methods (*13*) for these interferometric systems also impede the possibility of further reduction of fluorescence background and therefore, challenges further enhancement of the achievable resolution in 4Pi based single-molecule systems.

To overcome those difficulties, we introduced a method based on coherent pupil functions to allow accurate modeling of 4Pi-PSFs and developed an algorithm to extract the position information content at the theoretical information limit while, at the same time, dynamically compensates the temperature-induced cavity drift. The confluence of these new analytical methods improves both localization precision (1-4 fold depending on axial position of the emitter, **Fig. 4C**) and bias (2-6 fold’ reduction quantified from the s.t.d. of deviations in 2 µm axial range **Fig. 4D**) in all three dimensions compared to existing 4Pi-SMSN systems (*9, 11*) and at the same time extends imaging volume in the axial direction. Together with a novel 4Pi-compatible light-sheet illumination method reducing the fluorescence background by more than 5-fold (compared to epi configuration), we demonstrated the combined approach allowed further improving the achievable resolving power of 4Pi single-molecule localization microscopy.

**Fig 4.**
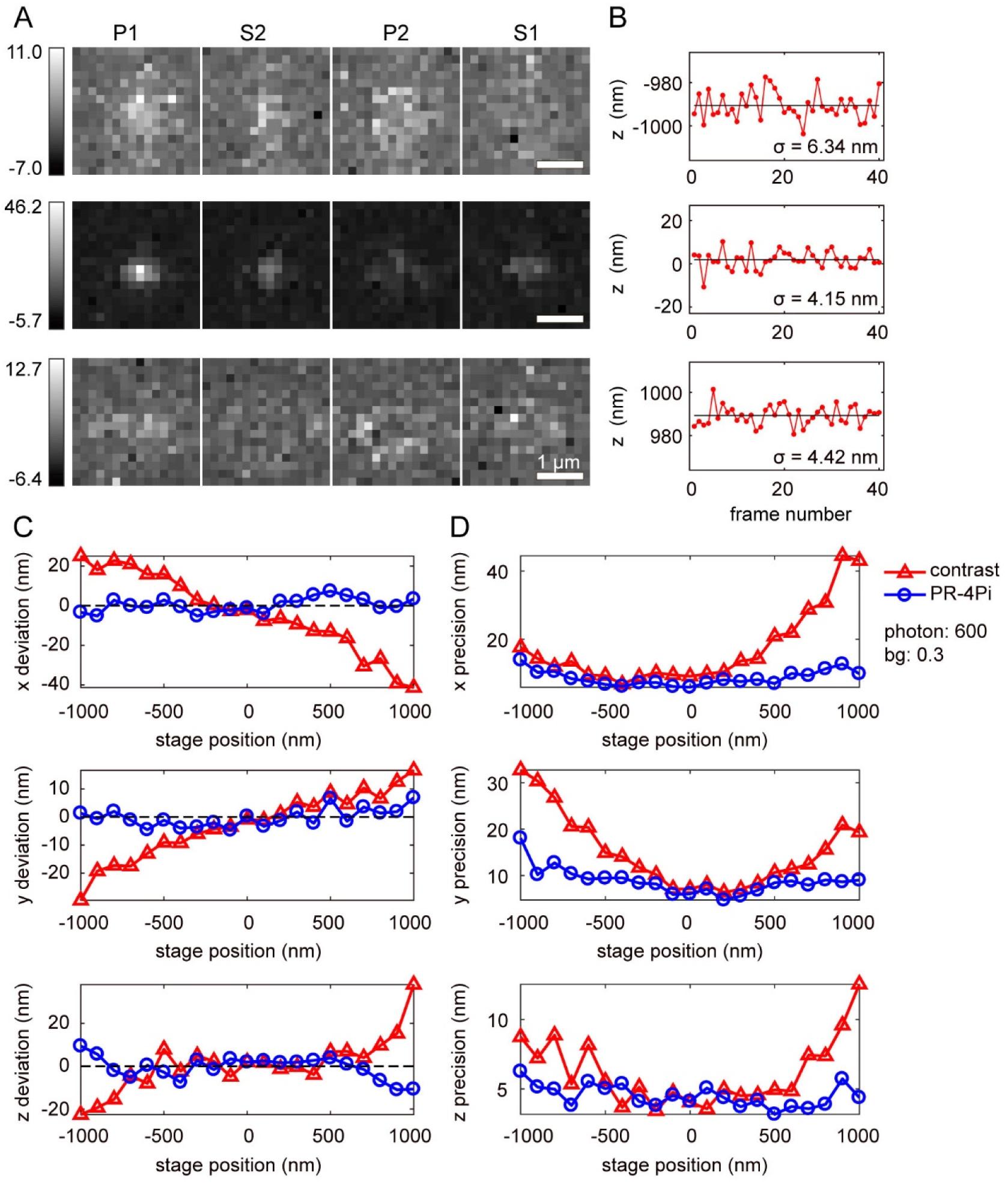
Comparison of localization results of bead data using PR-4Pi and contrast methods. (**A**) Examples of experimental bead images with relatively low detected photons at various axial positions in (B), each contains PSF patterns from four channels (P1, S2, P2 and S1). The pixel values represent the number of photon-electrons corrupted with sCMOS readout noise (*19*). (**B**) Axial localization results of the corresponding bead data in (A). Notice that in spite of the low photon budget, PR-4Pi localization method achieves a precision ∼5 nm. The estimation results fluctuated around the mean (solid black line) with a standard deviation of *σ*. (**C**) Localization deviations in *x, y* and *z*. (**D**) Localization precisions in *x, y* and *z*. The PR-4Pi method shows significant improvement in both localization accuracy (C) and precision (D). The bead data were acquired by imaging a fluorescence bead at axial positions from -1 to 1 µm by translating the sample stage with a step size of 100 nm, 40 frames were captured at each axial position (**Materials and Methods**). The estimation results from PR-4Pi method yielded a mean total photon of 600 per objective and a background photon of 0.3 per pixel. Contrast method in this work refers to the algorithm developed in (*11*) that extracts the axial positions using the Gaussian-weighted 0^th^ central moment (M0, **note S1** and **table S1**). **Figures. S3** and **S4** show comparisons with other 4Pi localization methods on the same bead data and data with higher photon count.

## Results

### Accurate 4Pi-PSF modeling with coherent phase-retrieved pupils

The hallmark of a 4Pi-PSF generated by an interferometric microscopy system is the distinct multi-lobe PSF in both axial and lateral dimensions. Currently, extracting single-molecule axial positions from the center lobes (including center peak intensity and intensities falls within a empirically defined ring region) has been the major focus in 4Pi single-molecule localization analysis (*8, 9, 11*), discarding significant amount of information carrying-photons among the lateral side lobes. At the same time, when pin-pointing molecular centers at the lateral dimension (i.e. localization in *x, y*), the interferometric features of the 4Pi-PSFs have been ignored (*8, 11*). As a result, no lateral resolution enhancement is reported comparing with conventional PSFs (*8, 11*).

To allow extracting information contained within the complex interferometric PSF, we describe here a PSF model based on two coherent pupil functions for 4Pi single-molecule imaging systems (*9, 11*). This method allowed us to accurately model the interferometric emission patterns taking into account of the independent wave front distortions from the two interference arms, the wavelength-dependent coherence modulation and the spontaneous optical path length drift.

Let *h*_*A*_ and *h*_*B*_ be a pair of pupil functions of the 4Pi system, representing the wave fields at the pupil planes of the upper and lower interference paths respectively (**Fig. 1**). The two pupil functions, retrieved independently through a phase retrieval algorithm (*14-16*), allowed us to incorporate aberrations for both interference paths. Interferometric PSFs were generated from the superposition of the wave fields, *h*_*A*_ and *h*_*B*_. Four superposed wave fields with different polarizations and phase shifts can be written as,

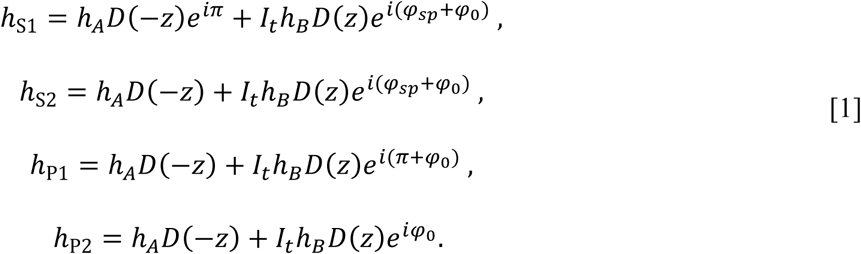

**Fig 1.**
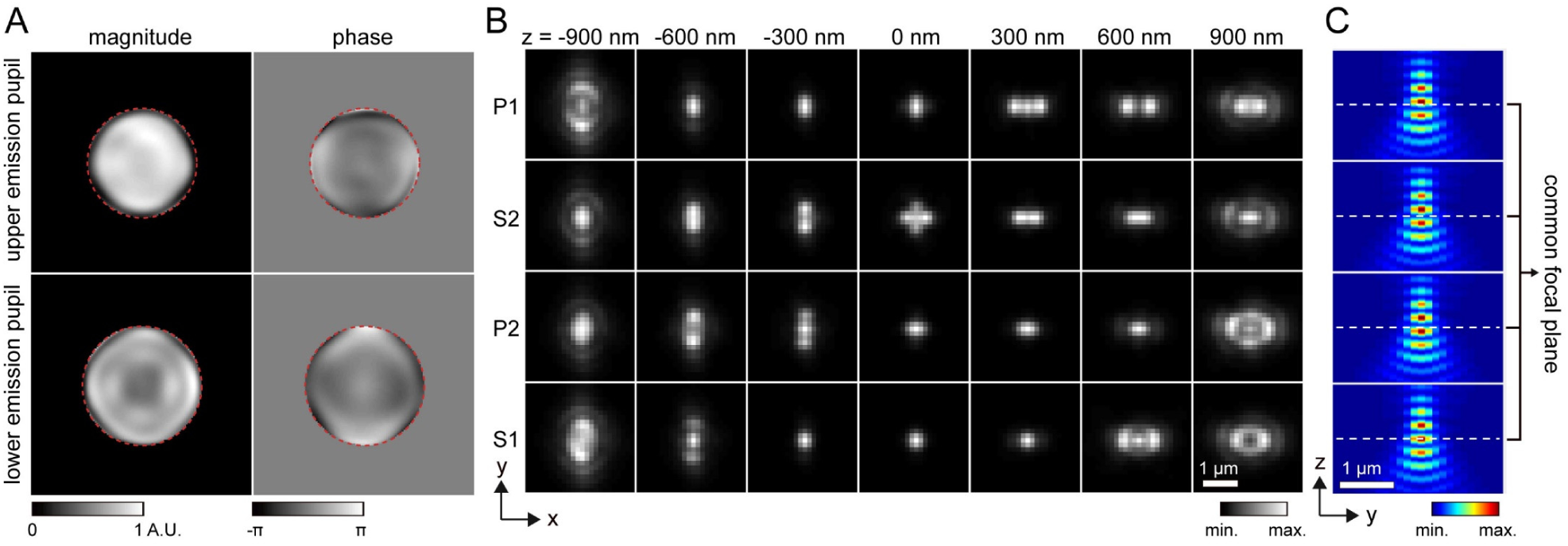
Coherent, phase retrieved pupil based 4Pi-PSF model (**A**) Pupil functions of upper and lower emission paths, independently retrieved by imaging a fluorescence bead on the bottom cover glass. Numerical apertures (NA) of 1.28 and 1.4 (the objective NA), defining the cutoff frequency in the Fourier space (red dash circles), were used during the phase retrieval of the upper and lower emission pupils. The slight shrinking of the upper emission-pupil size is caused by index mismatch aberration (**note S3**). Astigmatism aberrations (the 5^th^ Zernike polynomial, Wyant order) (*26*) with amplitudes of 1.5 and 2 (unit: 2πλ) were applied to the upper and lower deformable mirrors respectively (**Materials and Methods**). (**B**) PR-4PiPSF models at various axial positions. Each axial position including four PSF patterns with different polarizations and phases (channels P1, S2, P2 and S1). (**C**) PR-4PiPSF models in the y-z plane corresponding to the four channels in (B).

Here, we assume *h*_*A*_ and *h*_*B*_ are independent against polarization directions, where *φ*_*sp*_ *= 2πδL(n*_*s*_ *– n*_*p*_) / *λ*, with *n*_*s*_ and *n*_*p*_ the refractive indices of s-and p-polarized light in quartz, and *5L* the quartz thickness difference between the two interference paths (**note S2**). *φ*_0_ is the phase difference between the two interference arms when the single emitter is in the common focus of the two objectives, thereafter referred as the cavity phase. The defocus term *D(z)* equals to exp(*ik*_*z*_*z)*, with *k*_*z*_ being the *z* (axial) component of the wave vector and *z* is the axial position of the single emitter. A factor *I*_*t*_ was introduced to account for the transmission efficiency difference between the two interference paths. Here we assume *I*_*t*_ is one (**note S4** and **fig. S11**).

We can then write out the coherent PSF, *µ*_*Im*_, in terms of Fourier transform of each superposed wave field,

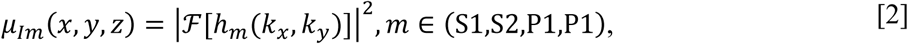

where subscript *I* indicates interference and Eq. **2** assumes that the wave fields *h*_*A*_ and *h*_*B*_ are perfectly coherent. However, because of the finite spectral width of the emission filter, the estimated coherence length of the emission light is small, ∼7.5 µm (with an emission filter of 700/50 nm, center wavelength/band width). Therefore, a slight change of the optical path length difference (OPD) between the two interference arms will result in a moderate reduction of the modulation depth – the peak to valley contrast in an interferometric PSF and the degree of this reduction is wavelength dependent. To this end, we assume *h*_*A*_ and *h*_*B*_ are partially coherent and the incoherent part will produce a conventional incoherent PSF described as *µ*_*W*_,

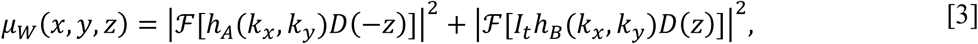

with subscript *W* indicates wide-field in the sense of conventional microscope. Therefore, the final PSF is a combination of the interferometric PSF and the conventional PSF with a factor *a* ∈ [0,1], describing the coherence strength (**note S4** and **fig. S11**),

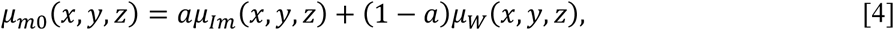

where *µ*_0_ represents the normalized PSF. By adding a total photon count of *I* and a background of *b*, the PR-4PiPSF model is,

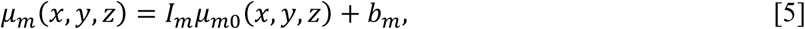

which includes quadruple PSFs for each emitter (**Fig. 1**). Here all *I*_*m*_ and *b*_*m*_ were considered as independent to account for the difference in transmission efficiency between the two polarizations and the emission paths after the beam splitter (*11*). We found that PR-4PiPSFs can produce relatively uniform resolutions within a large *a* range (0.5 to 1), an attractive feature for thicker specimens, where *a* is depth dependent (**fig. S11** and **note S4**).

As a demonstration of the accuracy of this 4Pi-PSF model, we compared it with the experimental PSFs obtained by imaging 100 nm beads attached on the coverslip surface. we found that the use of coherent, phase retrieved pupil functions can accurately model the experimental PSFs, while, in contrast, simulations of ideal 4Pi-PSFs (with unaberrated pupil functions) fails to represent the correct features (**Fig. 2**). This observation is further quantified through cross-correlations between different PSF models with the experimental PSFs. Comparing to the ideal-4PiPSF model which shows significant reduction of the correlation quantity at various axial positions, the developed method predicted an accurate PR-4PiPSF model showing strong agreement throughout a large axial range when compared to the experimental obtained dataset (**Fig. 2A** and **B**). We also compared the axial modulation of the PSF center intensity, we found that the PR-4PiPSF model, with a pre-calibrated coherence strength factor *a*, provides a good prediction on the modulation strength matching the measured interference PSFs obtained from a 100 nm fluorescent bead (**Fig. 2C**).

**Fig 2.**
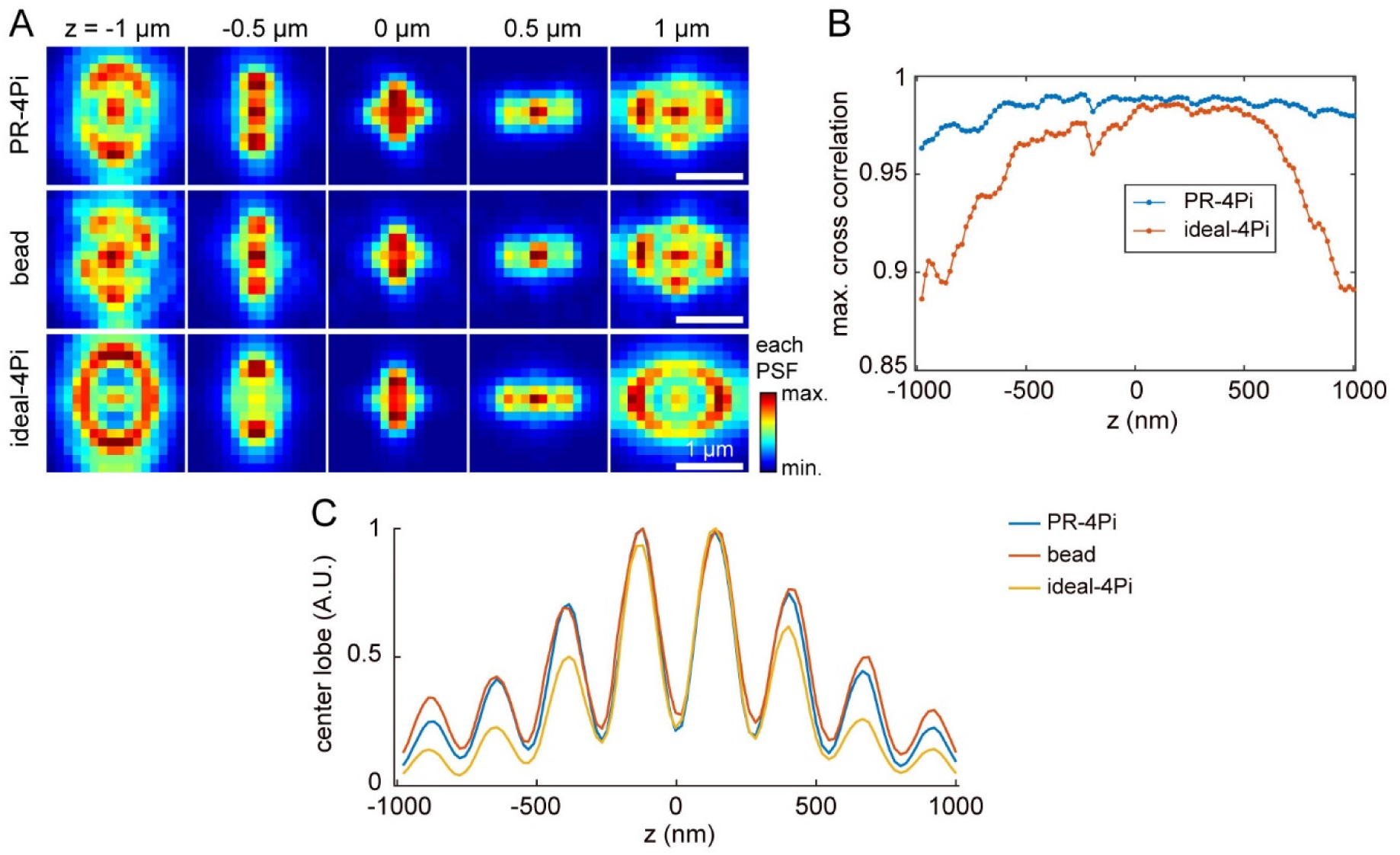
Comparison of phase retrieved 4Pi-PSF and ideal 4Pi-PSF with acquired 4Pi emission patterns from 100 nm bead. (**A**) PSFs at channel P2 from experimental bead data, phase retrieved (PR) and theoretical ideal PSF models. Ideal-4PiPSF model assumes no aberration, transmission loss and beam splitting inequality present in the imaging system. The simulated PSF models were generated based on the localization results (*x, y, z*) of the bead data using corresponding PSF models to find the best match in both PR-4Pi and ideal-4Pi cases. (**B**) Model accuracies quantified by cross correlations of PR and ideal 4Pi-PSFs with the experimentally acquired PSFs from beads. At each axial position, first the maximum cross correlation value between the PR/ideal-4PiPSF and the bead-image from each of the four channels (P1, S2, P2 and S1) was calculated, then the mean of the obtained four values was used to generate the curve. (**C**) Modulation intensity of the center lobes (Gaussian-weighted 0^th^ central moment) of various 4Pi-PSFs shown in (A) (**note S1**). The bead data were collected with high signal to noise ratio (SNR) at *z* positions from -1 to 1 µm, with a step size of 10 nm (**Materials and Methods**).

To further explore the effect on single-molecule localization with and without our PSF model, we pin-pointed 3D position of isolated beads with various photon counts and compared with the position readout from a high-precision piezo stage (±0.5 nm close-loop) (**fig. S3** and **S4**). We found that at nearly all the photon count levels, the lateral localizations of the beads at different axial positions almost completely overlap, while using an ideal PSF model results in increased deviations from the ground truth positions and worse localization precisions.

### Information content within an interferometric PSF

Given the ability to generate a realistic 4Pi-PSF model, we are able to quantitatively investigate the information content within the 4Pi-PSF pattern recorded on our microscope using Fisher information matrix (*17, 18*). By quantifying information content pixel by pixel within the 4Pi-PSF (as similarly shown previously (*10*) for ideal PSFs), we found previous 4PiSMSN localization analysis either discarded the additional information from interferometric detection (as for lateral localization, **note S8** includes derivation of a general case for such information gain) or incorporated only part of it (as for the axial localization that uses 0^th^ and/or 3^rd^ central moments of the PSFs, **fig. S1** and **S2**). Incorporating a comprehensive and accurate PSF model allows us to further improve both accuracy and precision in three dimensions.

To quantify the information content of 4Pi-PSFs, we calculated the Fisher information matrix, which quantifies the lower bound of the localization precision for an unbiased estimator: the Cramér-Rao lower bound (CRLB) (*18*),

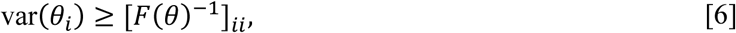

where *F* is the Fisher information matrix, *θ* is a vector of estimation parameters, *i* denotes the index of each parameter.

Incorporating Poisson noise and the readout noise from an sCMOS camera, a numerical calculation of each element in Fisher information matrix is (*19*),

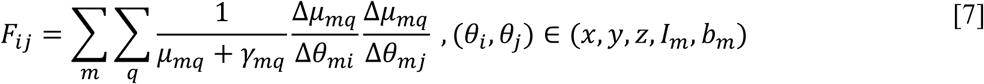

where *q* is the pixel index and *m* is the index of the quadruple 4Pi-PSFs, the factor *γ* equals to *σ*^2^*/g*^2^, where *σ*^2^ is the variance of the readout noise and *g* is the gain of each pixel.

In comparison with incoherent dual-objective PSFs (*20*), we observed the predicted information in a 4Pi-PSF increase significantly along the axial dimension (20-160 folds at 1000 total emitted photon and 10 background photons) and a relatively large information gain in the out of focus regions along the lateral dimension (4-7 folds at 1000 total emitted photon and 10 background photon) (**Fig. 3**).

**Fig 3.**
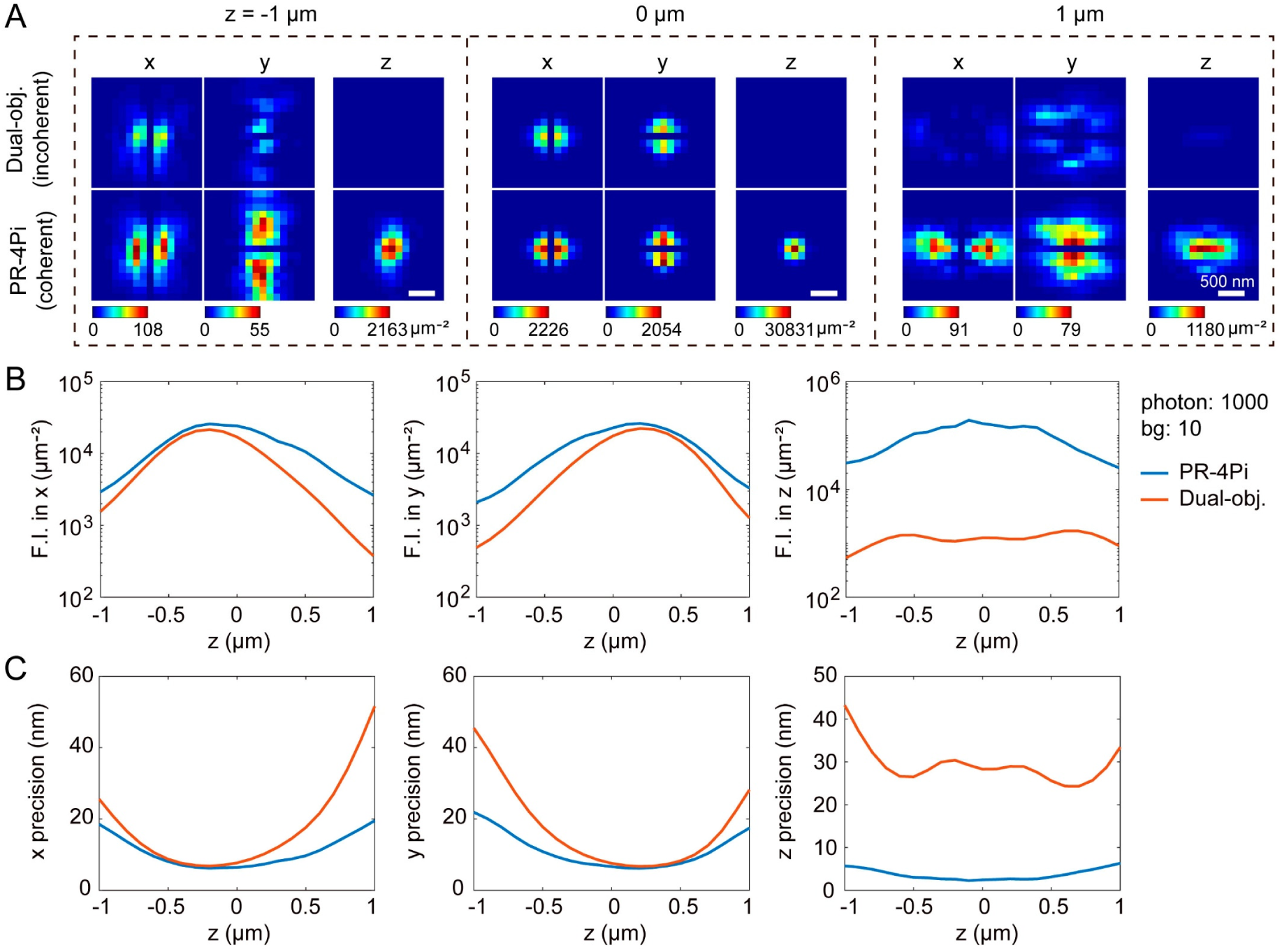
Information content comparison and theoretical localization precision limit of PR-4PiPSF and dual-objective (obj.) PSF. (**A**) Lateral distributions of the Fisher information (*17*) content per pixel in *x, y* and *z* estimations. Same parameters were used for both PR-4Pi and dual-objective configurations, except that the dual-objective configuration uses incoherent detection (**note S8**). (**B**) Fisher information (F.I.) content in *x, y* and *z* estimations. (**C**) Theoretical localization precisions in *x, y* and *z*. A total photon of 1000 per objective and a background (bg) photon of 10 per pixel were used for the PSF model generation. Phase retrieved pupil functions were used for both PR-4PiPSF and dual-objective PSF.

This observed information content boost predicts and quantifies the improvement in the achievable localization precisions in all three dimensions by using 4Pi-PSFs. Existing 4Pi localization methods obtain the lateral position estimation by either merging the multi-channel 4Pi-PSFs into incoherent dual-objective PSFs (*8, 11, 21*) or modifying the PSFs with an Gaussian mask (*9*) and applying a Gaussian or Airy disk PSF model on the merged or modified PSFs (**table S1**). These methods result in reduction of information since the interference pattern isn’t considered leading to up to 2-fold worse localization precisions when compared to the information limit, of which the deterioration is increasingly pronounced when emitters are located away from the common focus of the two objective lenses (**Fig. 3, fig. S1** and **S2**). As for axial localization, previously, position estimations were extracted from intensity contrast calculated from PSFs at different detection channels and such calculation often involves summing over all or partial pixel values (e.g. using the central moments, **table S1**) within the PSF of each channel. We found that these methods perform well within a small distance (±300 nm) from the common focus region but result in partial information loss, where the achieved precision (uncertainty value) and bias of these methods increase rapidly with the axial position of a single emitter (**fig. S1** and **S2**).

In order to utilize the complete information contained within the interferometric PSF pattern and minimize localization biases, we combined our 4Pi-PSFs model with a maximum likelihood estimator (MLE) with the appropriate noise model (*19*) (sCMOS noise model in our case) (**notes S1** and **S7)**. We demonstrated in the following section that the PR-4Pi algorithm achieve the theoretical precision limit (defined by CRLB) of our 4Pi-SMSN systems in both axial and lateral directions.

As a demonstration of PR-4Pi localization algorithm, first we tested the algorithm through a set of simulated 4Pi-PSFs (**Fig. 1**). While the localization precisions using conventional intensity-contrast based methods varies drastically in different axial position, we found PR-4Pi algorithm provides localization precisions consistently approaching the theoretical information limit calculated by CRLB. A detailed characterization of PR-4Pi algorithm and other existing 4Pi localization algorithms are shown in **fig. S2** (**table S1**). For fair comparison with existing 4Pi algorithms based on central moments for axial localizations (*9*), we generate 4Pi-PSFs without astigmatism modification (astigmatism modification were used in (*11, 21*) to reduce localization artifacts). We found that combining 0^th^ and 3^rd^ central moments (*9*) achieves better precision at out-of-focus region than using only the 0^th^ moment, but still fail to achieve the theoretical limit (CRLB) when considering equal split of photon energy through the four detected channels (5 estimation parameters include: *x, y, z, I* and *b*). Interestingly, when considering independent intensity and background for each channel (11 estimation parameters), the CRLB for axial estimation increases dramatically for near focus region, which results from an anti-correlation between estimations on intensity/background and axial positions without astigmatism modification (**fig. S1**). Nonetheless, PR-4Pi algorithm achieves CRLB in both parameter settings at nearly the entire tested ranges, except a small dip in the near focus region when considering independent intensity and background which is caused by a limited step size in the iterative optimization routine.

Further, we tested localizations on 100 nm bead data with photon counts in the range of experimental single-molecule emissions using fluorescent proteins such as mEOS3.2 (*22*) (∼600 photons per objective lens, 0.3 background photons). In consistent with our simulated results, PR-4Pi allows pin-pointing the position of a single bead with improved precision and bias than previous methods (**Fig. 4**). In addition, when assuming fixed intensity split between four quadrants, PR-4Pi results in improved axial precision, but at the sacrifice of increased biases (**fig. S3** and **S4**). Therefore, we propose to use PR-4Pi with independent background and intensity for each quadrant when minimizing localization bias is of most importance (**fig. S18**).

### Continuous cavity phase measurement

Another key feature in 4Pi/interferometric systems is cavity phase shift. The cavity phase describes the optical path length difference (OPD) between the two interference arms when the emitter is in focus at the common focal plane of the two objectives. The cavity phase, *φ*_0_, modulates with path length fluctuations and the changes of refractive indices of materials which photons travel through along the interference arms (approx. ∼600 mm each arm) and *φ*_0_, in turn, changes 4Pi-PSF from one pattern to another rapidly. We found the fluctuation of the cavity temperature lead to significant drift in *φ*_0_ (**note S5** and **fig. S5**). The cavity phase drift (i.e. *φ*_0_ drift) is different from sample drift. It is a unique challenge for interferometric systems. In comparison with commonly-observed sample axial drift which would make in-focused object out of focus, the cavity phase drift simply shifts the energy distribution between the four detected quadrants while an in-focus emitter remains in-focus. Here we examine the properties of cavity phase and propose a cavity-phase drift-correction algorithm.

The cavity phase drift, left uncorrected, degrades the axial resolution achievable in a 4Pi systems and creates image artifacts in the final reconstruction. We found a 0.04 rad drift per second (**fig. S5**) will cause noticeable axial-resolution deterioration in 10 s (equivalent to ∼18 nm axial drift, **fig. S6**), mandating a correction per 10 s to avoid resolution deterioration. This is especially challenging for the previous W-4PiSMSN method as the number of emitters collected within 10 s is not sufficient to correctly perform, ridge-finding, phase unwrapping and subsequent drift correction (*11*). In contrast, pupil-based method allows bypassing these problems by first no longer relying on phase unwrapping to recover *z* positions and second automatically compensating the cavity phase drift during the regression step (**note S1** and **fig. S7**).

We are able to obtain reliable calibration of *φ*_0_ from each 10 s data batch, so that a unique PR-4PiPSF model for a specific segment of time was generated to account for the *φ*_0_ drift. This constricted the axial resolution deterioration within 5 nm on average (**note S1**). In short, within a short segment of blinking dataset, total phase *φ* and shape metric *σ*_*s*_ of each emitter were measured (**notes S1** and **S6**). Assuming linear dependences of *φ* with the axial position of the molecule as well as the shape metric *σ*_*s*_ within a 2π range, the cavity phase *φ*_0_ was estimated as the total phase *φ* at the common focus plane of the two objectives. For single-section imaging, assuming a smooth change of the cavity phase during imaging, the estimated *φ*_0_ over time were interpolated with a smooth spline to eliminate abrupt transitions generated by estimation uncertainties and the interpolated values were incorporated into the PSF model during the regression step (**note S1** and **fig. S7**).

For data acquired at multiple optical sections for large volumetric 3D imaging, the scanning of the specimen to image different optical sections in the axial direction also changes the cavity phase. Assuming the scanning step size *d* is a constant during the data acquisition, this additional phase can be calculated from,

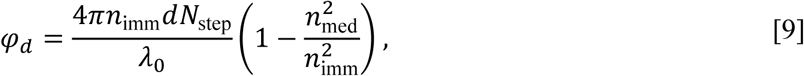

where *λ*_0_ is the emission wavelength in air, and *N*_step_ is the step number, which is indexed from 0. We would like to note that when the refractive index of the immersion media *n*_imm_ matches the one of the sample media *n*_med_, sample scanning in axial direction no longer causes cavity phase change. Eq. **9** allows us to quantitatively connect the cavity phases obtained at different optical sections and therefore, results in a continuous phase curve which improves the robustness of cavity phase calibration. Together, we are able to calibrate cavity phase based on short single-molecule blinking dataset (∼10 s) with high consistency across multiple optical sections throughout a cell. We found that for both single-section and multi-section imaging the calibrated curves match well with the estimated *φ*_0_ and are robust to the estimation noise (**note S1** and **fig. S7**).

We tested the reliability of our cavity phase measurement by localizing bead images at various *z* positions (**fig. S12**). An incorrect cavity phase can result in nearly 50% ghost images and when using an ideal-4PiPSF model, in spite of cavity phase values, the smallest ghost-image percentage is nearly 30% of the entire localizations. In contrast, we found that using a correctly retrieved cavity phase together with PR-4Pi algorithm, the percentage of ghost images is limited to 5-10% for both bead and cell imaging. For imaging in biological specimens, additional ghost reduction procedures further reduce it to ∼2% (**note S1, figs. S7** and **table S1**).

### Background reduction with highly inclined light-sheet illumination

In addition to accurately model PSF and track cavity phase, background fluorescence represents another key factor in determining the achievable resolution within a cell or tissue specimen. Here, we developed a light-sheet illumination modality that is compatible with the complex geometry of an interferometric imaging system allowing us to reduce the fluorescence background by more than 5-fold (**Fig. 6** and **fig. S8**).

**Fig 6.**
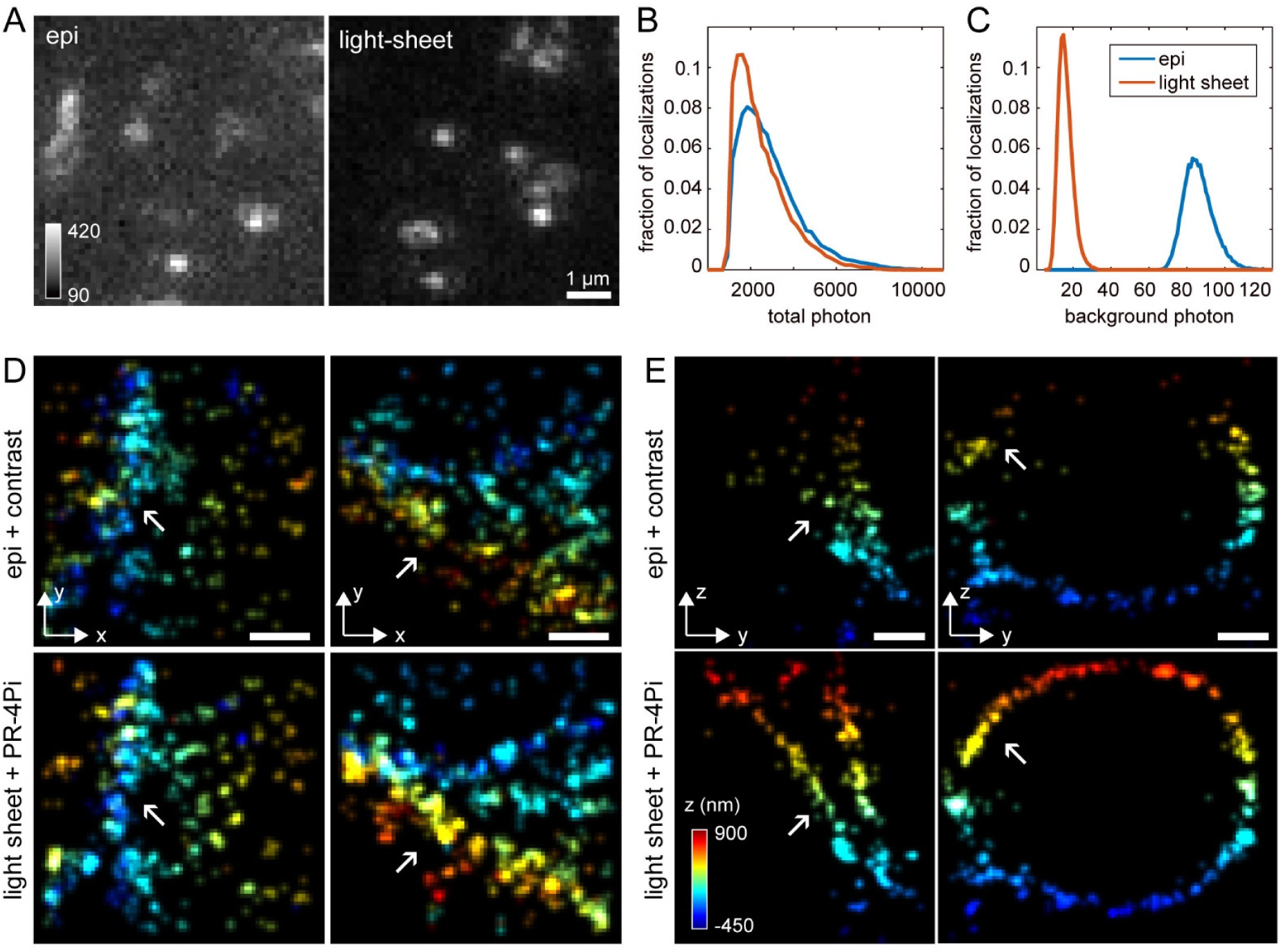
Background reduction and resolution improvement in both lateral and axial directions using light-sheet illumination and PR-4Pi localization method. (**A**) Example regions from raw blinking data under epi and light-sheet illuminations. Pixel values represent the direct readout values (ADUs, analog to digital unit) from a sCMOS camera (**Materials and Methods**). (**B, C**) Distributions of estimated total photon (from both objectives) and background photon (sum of the four channels) counts of single emitters from SMSN data acquired under epi and light-sheet illuminations. (**D, E**) SMSN reconstructions of immune-labeled TOM20 protein in COS-7 cell at the x-y and y-z planes. Data from epi illumination were analyzed with contrast method, while data from light-sheet illumination were analyzed with PR-4Pi method. The sample was imaged at a single axial plane with 20,000 frames under epi illumination followed by 20,000 frames under light-sheet illumination (**fig. S8**). Scale bars: 200 nm in (D, E).

Specifically, we design the excitation beam to split into two independent paths: an epi illumination from the upper objective and a highly inclined light-sheet (Hi-LS) illumination from the lower objective. The epi illumination was used for locating regions of interest and switching fluorophores to the dark state before single molecule imaging. The Hi-LS path generates a light sheet by using a cylindrical lens in the lower excitation path. The steering of the light sheet was accomplished by mirrors at the conjugate image and pupil planes, and the thickness of the light sheet was adjusted by a slit after the cylindrical lens. Comparing with the HILO illumination (*23*) that was generated by tilting a narrow epi illumination beam, Hi-LS tilts a light sheet resulting in a thinner section (an estimated thickness of 2.3 µm) and a larger field of view (FOV, ∼ 22 × 8 µm^2^. As for HILO illumination in our designed 4PiSMSN system, the sheet thickness will be 6-16 µm for an illumination area of 8-22 µm in diameter from the theoretical calculation, therefore, resulting in hardly any background reduction for 4PiSMSN systems) (**note S9** and **fig. S8**).

From raw single-molecule blinking data with epi and Hi-LS illumination, we found that the background is reduced significantly from 85 photons to 15 photons (at the peaks of the background photon distributions, **Fig. 6C**), a more than 5-fold reduction, which could improve the localization precisions by ∼1.4-1.7 fold (predicted by CRLB when using 11 estimation parameters, **fig. S16**). Further, we performed 4Pi-SMSN imaging on TOM20 in COS-7 cells comparing epi and Hi-LS illumination methods. Each reconstructed through 20,000 4Pi-SMSN blinking frames, the results show improvement in both lateral and axial resolutions using Hi-LS compared to epi. With the increased contrast, Hi-LS illumination also extends the localization range along the axial dimension (**Fig. 6D-E** and **fig. S10**).

We imaged immune-labeled TOM20 protein in COS-7 cells. For single optical-section data, a total of 160 distinct PR-4PiPSFs, each representing the concurrent state during a 10 s imaging period, were generated for localization analysis (**fig. S7**). For multi-section volumetric 4Pi-SMSN imaging, 240 distinct PR-4PiPSFs (10 s per PSF) were used to obtain three optical volumes spaced 800 nm apart (**fig. S7**). These time dependent 4Pi-PSFs allow capturing the concurrent states of the interferometric system with high accuracy and precision, and therefore enhance the resolution in both lateral and axial directions as demonstrated in the reconstruction of TOM20 clusters (**Fig. 5** and **fig. S9**). We found that with PR-4Pi algorithm, those clusters appear to be brighter and distinct, indicating higher localization precision and acceptance percentage. The capability of localizing more emitters or emission events will benefit subsequent processing such as particle averaging and emission-event combining to achieve even higher resolution. In addition, together with extended axial localization range, PR-4Pi achieves higher alignment precision of optical sections for volumetric imaging (**fig. S17**).

**Fig 5.**
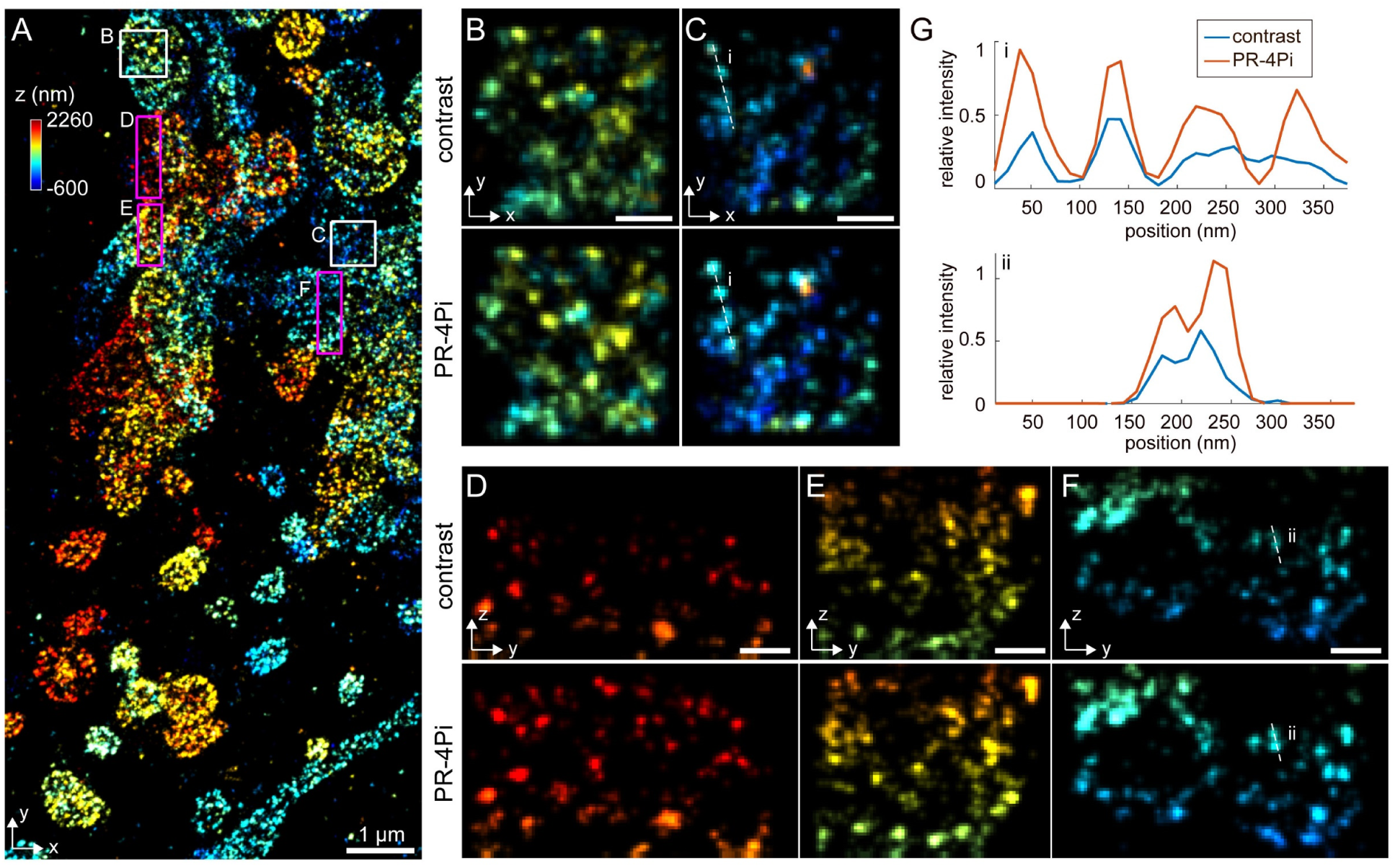
SMSN reconstruction of a 3D volume of mitochondria (TOM20) in COS-7 cells. (**A**) Lateral view of the SMSN reconstruction with each localization color-coded by its axial position. (**B, C**) Zoom-in regions in (A, white boxes) at the x-y plane demonstrating constriction of the TOM20 cluster size using PR-4Pi localization method. (**D-F**) Zoom-in regions in (A, magenta boxes) at the y-z plane. (**G**) Intensity profiles along the dotted lines in (C) and (F), scaled by the maximum intensity of the selected regions from the PR-4Pi method. Color scaled with the axial position. Scale bars: 1 µm in (A), 200 nm in (B-F).

Furthermore, by combining the developed light sheet illumination with PR-4Pi localization algorithm, the capacities of resolution enhancement from both methods are multiplied (**Fig. 6D** and **E**). We found that the visualization of the cross-sections in both axial and lateral directions show tighter and brighter TOM20 clusters demonstrating the achieved higher resolution in all three dimensions (**Fig. 6D** and **E**). Line profiles across multiple clusters confirmed the observation and shows clearly separated cluster peaks while contrast method fail to resolve these ultra-structures (**Fig. 5G**). Furthermore, a full axial range of 1.4 µm per optical section can now be recovered (**Figs. 5D** and **6E**, and **fig. S9**).

## Conclusion

In this work, we developed a coherent, phase-retrieved pupil based single-molecule localization algorithm for 4Pi-SMSN systems. Investigating the 4Pi-PSF information content within each pixel revealed that PR-4Pi localization algorithm is capable of achieving the estimation lower bound predicted by Fisher information content.

Comparing with existing 4Pi single-molecule localization algorithms, we demonstrated that PR-4Pi allows further improvement on both precision and bias in all three dimensions. In addition, we developed a 4Pi compatible light-sheet illumination method using a highly inclined light sheet. This method significantly reduces the fluorescence background by more than 5-fold compared to epi illumination. By combining the novel localization algorithm and light-sheet illumination for 4Pi/interferometry based single-molecule super-resolution system, we demonstrate the possibility of further pushing the boundary of achievable 3D resolution in interferometric single-molecule imaging systems.

## Materials and Methods

### Sample preparation

Before SMSN imaging, round coverslip (25 mm diameter) containing immune-stained COS-7 cells was incubated with 200 µL bead dilution, containing 100 nm crimson bead (custom-designed, Invitrogen) diluted to 1:10^6^ in PBS, for 10 minutes. Then the coverslip was washed three times with PBS and placed on a custom-made sample holder. Then 150 µL imaging buffer (10% (w/v) glucose in 50 mM Tris (JT4109-02, VWR), 50 mM NaCl (S271-500, Fisher Scientific), 10 mM MEA (M6500-25G, Sigma-Aldrich), 50 mM BME (M3148-25ML, Sigma-Aldrich), 2 mM COT (138924-1G, Sigma-Aldrich), 2.5 mM PCA (37580-25G-F, Sigma-Aldrich) and 50 nM PCD (P8279-25UN, Sigma-Aldrich), pH 8.0) was added on top of the coverslip. Then a cleaned coverslip of the same size was carefully placed on top of it and the excessive buffer was removed with Kimwipes. The sample was sealed with two-component silicone sealant (Picodent Twinsil, Picodent, Germany).

For beads imaging, a cleaned round coverslip (25 mm diameter) was incubated with 200 µL bead dilution, containing 100 nm crimson bead (custom-designed, Invitrogen) diluted to 1:10^6^ in PBS, for 10 minutes. Then the sample was rinsed with PBS, drained and placed on a custom-made sample holder. Then 10 µL PBS was added on top of the coverslip and a second pre-cleaned coverslip was placed on top and the sample was sealed with two-component silicone sealant (Picodent Twinsil, Picodent, Germany).

For light-sheet profile measurement, the dye coated coverslip sample was prepared by first adding 200 µL poly-L-lysine (P4707-50ML, Sigma-Aldrich) onto a 25 mm round coverslip and was incubated for 1 hour. Then the sample was rinsed once with deionized water and was incubated with 200 µL 1:10^6^ dye dilution in 0.1 M sodium bicarbonate (792519-500G, Sigma-Aldrich) for 2 hours. The dye dilution was prepared from a stock solution that was made by dissolving Alexa Fluor 647 (A20006, Life Technologies) powder in DMSO (276855-100ML, Sigma-Aldrich), the color was dark blue. The sample was then rinsed three times with deionized water and mounted on a custom-made sample holder. Then 20 µL PBS was added on top of the coverslip and a second pre-cleaned coverslip was placed on top and the sample was sealed with two-component silicone sealant (Picodent Twinsil, Picodent, Germany).

### Immunofluorescence labeling

COS-7 cells (CRL-1651, ATCC) were seeded on 25 mm diameter coverslips (CSHP-No1.5-25, Bioscience Tools, San Diego, CA) 1∼2 days before immunofluorescence labeling. Cells were first rinsed three times with pre-warmed (at 37 °C) phosphate buffered saline (PBS, 806552-500ML, Sigma-Aldrich) and then fixed for 15 minutes at room temperature (RT) with pre-warmed (at 37 °C) 3% paraformaldehyde (PFA, 15710, Electron Microscopy Sciences, Hatfield, PA) and 0.1% glutaraldehyde (GA, Electron Microscopy Sciences, 16019, Hatfield, PA) in PBS. Cells were then washed twice with PBS and treated for 7 minutes in freshly-prepared fluorescence quenching buffer (0.1% sodium borohydride (452882-25G, Sigma-Aldrich) in PBS). After fluorescence quenching, cells were washed three times with PBS and treated for 10 minutes with 10 mM Tris (pH 7.3, JT4109-02, VWR). Cells were then rinsed three times with PBS and permeabilized with blocking buffer (3% bovine serum albumin (BSA, 001-000-162, Jackson ImmunoResearch) and 0.2% Triton X-100 (X100, Sigma-Aldrich) in PBS) for 30 minutes, gently rocking at RT. After blocking, cells were incubated with anti-TOMM20 primary antibody (sc-11415, Santa Cruz Biotechnology), diluted to 1:1000 in 1% BSA and 0.2% Triton X-100 in PBS, at RT for 1 hours. Cells were then washed three times each time for 5 minutes with wash buffer (0.05% Triton X-100 in PBS) and incubated with secondary antibody conjugated with Alexa Fluor 647 (A21245, Life Technologies, Grand Island, NY), diluted to 1:1000 in 1% BSA and 0.2% Triton X-100 in PBS, at RT for 1 hours. After incubation with secondary antibody, cells were washed three times each time for 5 minutes with wash buffer. And then cells were post-fixed with 4% PFA in PBS for 10 minutes. After post-fixation, cells were rinsed three times with PBS and stored in PBS at 4 °C until they were imaged.

### Data acquisition

All data were acquired on a custom-built 4Pi-SMSN microscope, constructed from the previous design (*11*) with implementation of Hi-LS or epi illumination (**fig. S8**). In Short, emitted photons were collected through two opposing high-NA oil immersion objectives (Olympus UPLSAPO 100XO, 1.4 NA) and received by a sCMOS camera (Orca-Flash4.0v2, Hamamastu, Japan).

The SMSN data were collected with the following procedure: 1) Optimize the upper and lower deformable mirrors (DM, MultiDM-5.5, Boston Micromachines) by imaging a fluorescence bead on the bottom coverslip. The optimization was accomplished by individually optimizing the five dominant mirror modes (*24, 25*), representing normal and diagonal astigmatism, horizontal and vertical coma and spherical aberration. The index mismatch aberration from the upper objective imaging through the sample medium was also compensated during this step (**note S3** and **fig. S13**). 2) Apply astigmatism to each DM, amplitudes of 1.5 and 2 (unit: 2πλ) for the 5^th^ Zernike polynomials (Wyant order) (*26*) were used for the upper and lower DMs respectively. 3) Collect a stack of incoherent PSF images through the upper and lower objectives independently by imaging a bead on the bottom coverslip at *z* positions from -1 to 1 µm, with a step size of 100 nm, a frame rate of 1 Hz and taking two frames per axial position. Those PSF data were used to generate phase retrieved pupil functions of upper and lower emission paths. 4) Align the *x, y* and *z* positions of both objectives by centering and focusing the beads images from those objectives, and then collect a stack of interference PSF images by imaging a bead on the bottom coverslip at *z* positions from -1 to 1 µm, with a step size of 10 nm, a frame rate of 1 Hz and taking one frame per axial position. This PSF dataset was used for the estimation of the quartz induced phase shift Δ*φ*_sp_ (**note S2**). 5) Focus on a cell that took up greater than 70% of the field of view and collect 40 frames of cell images through only the lower objective at a frame rate of 10 Hz. This cell data was used for quadrant alignment (**note S6**). Steps 1 to 5 must be repeated for each sample to ensure a realistic modeling of the 4Pi-PSF and an accurate alignment of the four quadrants. 6) Locate a region of interest (ROI) and realign the objectives. Steps 1 to 6 were performed under epi illumination with a 642 nm laser (2RU-VFL-P-2000-642-B1R, MPB Communications Inc., Canada) at an excitation intensity of 90 W/cm^2^. 7) Briefly illuminate the ROI with a laser intensity of 7∼9 kW/cm^2^ under epi illumination then switch to Hi-LS illumination. During data acquisition, a 405 nm laser (DL-405-100, CrystaLaser) was used as an activation laser and was gradually adjusted from 0 to 20 W/cm^2^. For single-plane imaging, 20,000 to 80,000 frames were collected. For multi-plane imaging, the cell was scanned axially by moving the stage at a step size of 800 nm from the bottom to the top of the cell for 10-20 cycles, 3-4 axial planes were imaged per cycle and 2000 frames were collected per axial plane in each cycle.

Bead data for testing PR-4Pi algorithm were collected by imaging a bead on the bottom coverslip at *z* positions from -1 to 1 µm, with a step size of 100 nm, and taking 40 frames per axial position. The bead intensity of each dataset was adjusted by varying the excitation laser power and the exposure time.

### Fisher information content of contrast method

Contrast method in this work refers to the algorithm developed in (*11*) that extracts the axial positions using the Gaussian-weighted 0^th^ central moment (M0, **note S1.3** and **table S1**). In the lateral dimension, the Fisher information content of contrast method (*11*) (**table S1**) is equivalent to the one using astigmatism method on a dual-objective system (*20*) (**Fig. 3**). In the axial dimension, the contrast method operates on the center lobe of each PSF, which was extracted by multiplying the PSF with a 2D Gaussian at a width of one pixel. After multiplication, the remaining information for axial localization is 20-60%.

## Acknowledgement

We thank E. Allgeyer and G. Sirinakis from University of Cambridge and Y. Zhang from Yale University for valuable suggestions on construction of 4Pi-SMSN microscope. We thank F. Xu for helpful discussions on algorithm development and suggestions to the manuscript. We thank D. A. Miller and B. Chen for suggestions on cell culture. S.L. and F.H. were supported by grants from NIH (R35 GM119785) and DARPA (D16AP00093). Data that support the findings of this study are available from the corresponding author upon request.

## Author Contributions

S.L. and F.H. developed the algorithm. S.L. wrote the software. S.L. performed the experiments and analyzed the data. S.L. and F.H. constructed the imaging systems. S.L. and F.H. conceived the study and wrote the manuscript.

## Code Availability Statement

MATLAB scripts including all functions, such as cavity phase calibration, ghost reduction, MLE localization and etc., associated with the PR-4Pi algorithm and an example script on using PR-4Pi algorithm will be made available upon publication as Supplementary Software and further updates will be made available at https://github.com/HuanglabPurdue/PR-4Pi.

## Supplementary Materials for

## Supplementary Notes

### 1. PR-4Pi localization algorithm

The PR-4Pi localization algorithm consists of four steps: calibration of cavity phase *φ*_0_, regression, drift correction (*1*) and ghost reduction.

#### 1.1 Calibration of cavity phase

First, the obtained blinking dataset from the camera were segmented into short time series (data batches) each contains 500 frames (10 s). For each data batch, cavity phase *φ*_0_ was obtained as described in the next section. From these time-dependent *φ*_0_ measurements, a continuous cavity phase function, *φ*_0_*(t)*, was obtained through interpolation which was used in generating the time-dependent PR-4PiPSF model at any specific data acquisition period and therefore compensates the *φ*_0_ drift with high accuracy.

The detailed calibration procedure is as follows: For every data batch (500 frames), 1) Sub-regions of 16 × 16 pixels around the centers of isolated emitters in each frame were cropped (*2*) after quadrant alignment (**note S6**). Each isolated emitter has four sub-quadrants; 2) Sum of the four sub-quadrants generates a conventional astigmatism PSF from which PSF widths *σ*_*x*_ and *σ*_*y*_ were estimated through a 2D Gaussian fitting (*3, 4*); 3) The four sub-quadrants of an isolated emitter were then used to find the interference phase *φ*, which includes both cavity phase *φ*_0_ and the phase shift introduced by emitter’s axial position within the specimen (**note S1.3**); 4) From a scatter plot of *φ* vs. *σ*_*s*_ obtained from a large number of isolated emitters within the data batch, where 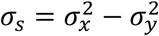, a single valued *φ*_0_ was estimated for the specific data batch (**note S1.2** and **fig. S7**). The entire dataset may contain up to 200 data batches each contains its own estimated *φ*_0_. This time-dependent sequence of *φ*_0_ values were then unwrapped and interpolated using a smooth spline method through MATLAB (MathWorks, Natick, MA) function *fit* (**fig. S7**). The obtained spline function represents the time-dependent cavity phase drift during the data acquisition and was used to generate a time-dependent PSF model during the regression step.

For volumetric imaging including multiple optical sections, the calibration process also need to consider the phase changes introduced by axial positions of optical sections, denoted as *φ*_*d*_. Assuming the step size between two optical sections is a constant during the data acquisition and denoted as *d*, then *φ*_*d*_ can be calculated from,

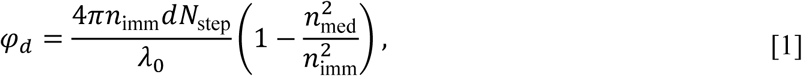

where *λ*_0_is the emission wavelength in air, and *N*_step_ is the step number, which is indexed from 0. The estimated *φ*_0_ for each axial plane were subtracted by the corresponding *φ*_*d*_ and the resulting phases again form a continuous curve as in the single-plane imaging and were fitted with a smooth spline to reduce the estimation noise. The generated spline curve represents the cavity phase evolution throughout the acquisition period excluding the extra phase difference introduced by optical sections’ axial positions. By adding back the expected *φ*_*d*_ for each optical section to above obtained spline curve, each optical section then carried its own time-dependent cavity phase evolution curve (**fig. S7**). Note that the *φ*_0_ were obtained based on a precalibrated *σ*_*s*0_ - the shape metric value *σ*_*s*_ when a PSF is in focus, obtained from imaging beads on the cover glass (**Materials and Methods**). But with index mismatch aberration, the width of the in focus PSF changes as the imaging depth increases. Therefore, the use of a constant *σ*_*s*0_ for different axial planes will result in an inaccurate estimation of *φ*_0_. However, for the demonstrations in this work, the maximum imaging depth is ∼3 µm, within which the effect of index mismatch aberration can be ignored (**note S3**).

With calibrated cavity phase evolution (*φ*_0calib_), cavity-phase induced axial-drift 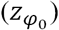 were automatically compensated every 10 s, within which an average of 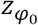 can be estimated from mean(*z*_rate_Δ*φ*_0calib_/2), where *z*_rate_ is the axial drift caused by 1 rad of cavity-phase drift and is 45 nm from simulation. We found that the averaged 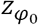 within 10 s was less than 5 nm for the demonstrations on mitochondria imaging (**Figs. 5** and **6, fig. S9** and **S10**).

#### 1.2 Estimation of cavity phase

The cavity phase *φ*_0_ for a data batch (**note S1.1**) was estimated as follows: due to wrapping of the estimated interference phase *φ*, the scatter plot *φ*-*σ*_*s*_ resembles multiple stripe patterns that repeats from 0 to 2π (**fig. S7**). These stripes were fit with multiple lines sharing a common slope. Subsequently, the linear fit along the stripe with the highest localization density was selected from the cross-correlation of a reference image and the Gaussian blurred *φ*-*σ*_*s*_ image along the *σ*_*s*_ dimension (**fig. S7**). Then, the selected linear fit was used to extract the cavity phase *φ*_0_ at *σ*_*s*_ *= σ*_*s*0_. The value of *σ*_*s*0_ was the mean of *σ*_*s*0_ estimates from five bead datasets (**fig. S7**). With a small standard deviation on *σ*_*s*0_ estimation, a single *σ*_*s*0_ value was used for all datasets (-0.3 pixel^2^ in our experiments).

#### 1.3 Regression

The regression step is to obtain the estimation parameters such as 3D locations, total photons and background photons of single emitters. It operated on small batches of the acquired data sequence. The batch size, which we used is 500 frames, is determined by the duration in which the cavity phase is relatively stable (**note S1.1**). A particular *φ*_0_ obtained from time-dependent cavity-phase calibration was used for each data batch (**note S1.1**).

First, the obtained four quadrants (from channels S1, S2, P1 and P2) in each camera frame were aligned and summed. The summation reduces the 4Pi emission patterns into conventional incoherent emission patterns and the summation images were used for 2D Gaussian fitting or astigmatism localization based on a Gaussian PSF model (*3*). Subsequently, from the summed image, subregions containing isolated single molecule emission events were cropped (*2*) and then were fitted with a 2D Gaussian to obtain the initial guesses of lateral positions (*x* and *y*), and the estimations of the PSF widths *σ*_*x*_ and *σ*_*y*_ (*3*). For initial estimations of the total photon and the background photon counts, the subregions were cropped without the summation of the four quadrants, so that each isolated single-molecule emission event contains four sub-quadrants. The background, *b*_*m*_, for each sub-quadrants, was estimated from the median value of all edge pixels in each sub-quadrant, and the total photon count, *I*_*m*_, was the sum of all pixel values for each sub-quadrant after background (*b*_*m*_) subtraction, where *m* denotes the four channels and *m* ∈ (S1,S2,P1,P2).

Second, emission patterns which have low interference contrast or introduce large inconsistency between conventional astigmatism and PR-4Pi estimations were removed by the following procedure: 1) Modulation contrast values were calculated for each emitter. To this end, we first computed the Gaussian-weighted sum of each sub-quadrant (*5*), denoted as *G*_*m*_,*m* ∈ (S1,S2,P1,P2), which is the sum of all pixel values of each sub-quadrant after multiplication of a 2D Gaussian with a width of one pixel (Gaussian-weighted 0^th^ moment (M0) as defined in (*5*), **fig. S1** and **S2**). Then the modulation contrast was calculated from (*1*),

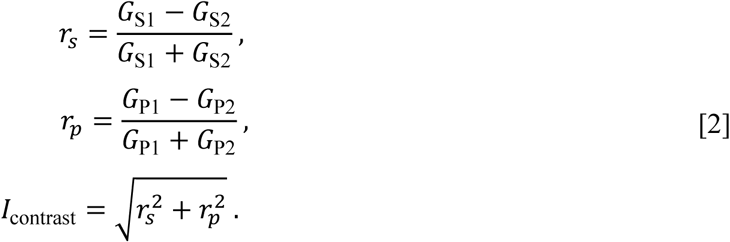

And emitters with a modulation contrast lower than 0.3 were discarded, since they usually contain overlapping emitters or have low signal to noise ratios (SNR). 2) Interference phase (*1*) *φ = arctan*(*r*_*p*_*/r*_*s*_) and the quantity 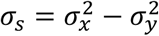 were calculated for each emitter. Emission patterns were discarded when the pair (*σ*_*s*_, *φ*) has a minimum vertical distance (*φ*_dis_) greater than 2π/3 to the linear fitted lines described in (**note S1.2** and **fig. S7**), representing large discrepancy in axial position estimations from two independent 3D modalities, i.e. astigmatism (*σ*_*s*_) and contrast based 4Pi (*φ)*, where *σ*_*s*_ and *φ* are linear with respect to axial positions. Theoretically, when noise and aberration are absent, all (*σ*_*s*_, *φ*) pairs should lay on top of the linear fitted lines (**fig. S7**).

Third, we seek to find reasonable initial guesses of axial positions from the remaining emission patterns. To this end, a stack of 51 PR-4PiPSFs at *z* positions from -1 to 1 µm with a given *φ*_0_ was generated and subsequently, initial axial position guesses were obtained through cross-correlations between each emitter’s sub-quadrants with the pre-obtained 4Pi-PSF stack (**fig. S7**).

At last, using the above obtained initial guesses on lateral and axial positions, total photon and background photon counts, a maximum likelihood estimator (MLE) was used to obtain estimates for all 11 parameters (referred as *θ*) (*6*). The likelihood function for each 4Pi emission pattern can be written as,

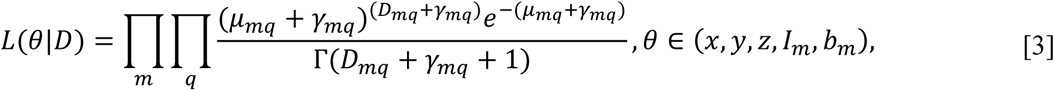

which includes the Poisson noise and pixel-dependent sCMOS noise (*6*). *D* represents the experimental data, *q* is the pixel index and *γ* equals to *σ*^2^*/g*^2^ where *σ*^2^ is the variance of the readout noise of each pixel and *g* is the gain of each pixel. Note that the subregions (*D*) used for MLE were cropped from original data quadrants without transformation (**note S6**). An optimization routine based on a modified Levenberg-Marquadt method (*7*) was used to minimize the negative log-likelihood function, which is given by, using Sterling approximation,

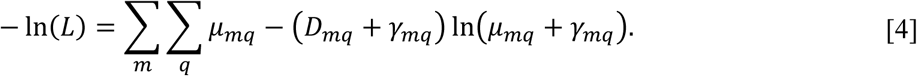

Its first and second derivatives are

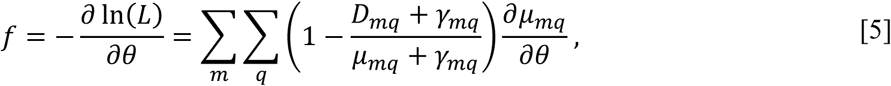

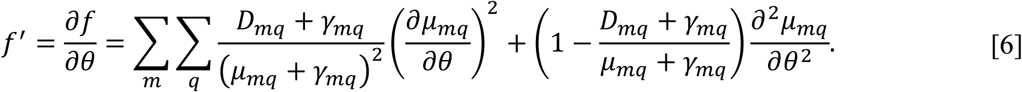

Here, we set the term with *∂*^2^*µ*_*mq*_*/ ∂ θ*^2^ in Eq. **6** to zero (*7*). Then the estimation parameters were updated from,

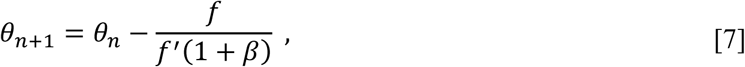

where *β* is a damping factor that was used to adjust the convergence speed and performance, here we set *β* to 1.

#### 1.4 Ghost reduction

The likelihood function used for regression has many local minimums along the axial dimension, which come from the periodicity in the intensity modulation of the interferometric PSFs. Those local minimums are separated by a distance close to the modulation period, ∼300 nm. With low SNR or overlapping emitters, the optimization process can be trapped at the local minimums and results in ghost images of a single emitter (*5, 8*). Here we developed a ghost reduction (GR) algorithm exploiting the periodicities of these local minimums to reduce the ghost images, which was applied both during and after the regression step (**fig. S7**). The GR algorithm consists of three steps: select a candidate image of the single emitter, calculate the ghost probabilities, update (during regression) or remove (after regression) the determined ghost images.

##### Selection of candidate image

1) Within each data batch of *N* frames, we calculate the lateral and axial pairwise distances *d*_*xy*_ and *d*_*z*_ for all localized emitters. 2) Emitters that are close in lateral dimension (*d*_*xy*_< 50 nm) but with an axial separation (270 < *d*_*z*_ < 320 nm) near the modulation period are grouped together as ghost pairs. Each ghost pair can contain multiple clusters of points, which are mutual ghost images of each other (**fig. S7**). 3) Select one cluster from each ghost pair as the candidate image of the single emitter based on a selection criterion,

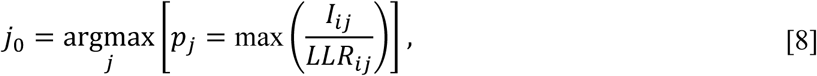

where *I* is the sum of estimated photons (*I*_*m*_, **Main Text** Eq. 5) of the four sub-quadrants, *j* the cluster index and *i* the index of each localization in the cluster. And *LLR* represents the log-likelihood ratio (*2, 9*) of each localization,

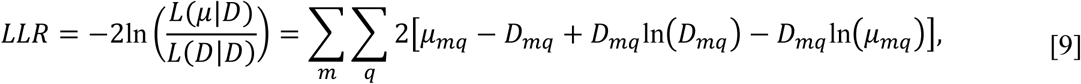

Where *q* is the pixel index and *m* is the channel index and *m* ∈ (S1,S2,P1,P2). The selection method first finds the maximum ratio of *I/LLR* within each cluster, denoted as *p*, then select the cluster with the largest *p* as the candidate image (cluster *j*_0_).

##### Calculation of ghost probability

We assume the candidate image is the real image of the emitter, a naive GR algorithm is to remove all the rest images of that emitter. Apparently, this simple method will remove any periodic localizations, with a period near the modulation period (*z*_*T*_), along the axial dimension and prone to change the sample structures. In fact, not all images in one ghost pair belong to a single emitter, some might be the real images of other emitters that happen to be ∼*z*_*T*_ distance away. Therefore, we calculated the probability of each cluster being a ghost image of the candidate cluster, named as the ghost probability. The detailed calculation is as follows: 1) For the same data batch, obtain axial position estimations using the conventional astigmatism localization method (*3*), denoted as *z*_ast_. The astigmatism method is ghost free, so that the real ghost localizations from PR-4Pi should collapse into one cluster when replacing the axial positions with *z*_ast_. 2) For each ghost pair, calculate the standard deviations of the estimations in *x, y* and *z*_ast_ of the candidate cluster, denoted as *σ*_*gx*_, *σ*_*gy*_ and *σ*_*gz*_, but restricted to a set range (10-30 nm for *x* and *y* and 40-100 nm for *z*_ast_). 3) Calculate the center of the candidate cluster from the medians of the estimations in *x, y* and *z*_ast_, denoted as *x*_0_, *y*_0_ and *z*_ast0_. 4) Calculate the centers of the rest clusters in the same ghost pair, denoted as *(x*_*i*_, *y*_*i*_, *z*_ast*i*_), then the normalized probability of cluster *i* belonging to the candidate cluster is

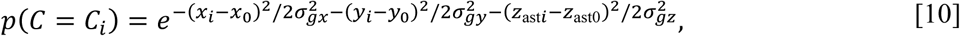

And the axial component of the above probability is

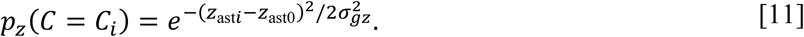

During regression, if either *p* > 0.001 or *p*_*z*_ > 0.1, we consider cluster *i* is a ghost image of the candidate cluster and set all the axial localizations in this cluster to *z*_0_, which is the median of the axial localizations of the candidate cluster from PR-4Pi. Otherwise, we consider cluster *i* as the localizations of a different emitter and keep the original localization results. 5) Assign the updated axial localizations as the new initial guesses of the *z* positions, and relocalize the emitters with PR-4Pi.

After regression, clusters with *p* > 0.001 or *p*_*z*_ > 0.1 will be removed. The data batch size *N* for GR during regression is 500 frames and 2000-20,000 frames for GR after regression. The two-step GR algorithm decreases the percentage of ghost images to 2% and the resulting distributions of pairwise distances are comparable with the ones from astigmatism method (**fig. S7**).

### 2. Estimations of hyper parameters in PR-4PiPSF model

In PR-4PiPSF model, besides the cavity phase *φ*_0_ and the Zernike coefficients (generated from the phase retrieval process), which are used to form the shape of the pupil function, there are a few hyper parameters are also crucial to build an accurate 4Pi-PSF model. Those hyper parameters are constants during the localization process and include: the refractive indices of the immersion medium *n*_imm_ and the sample medium *n*_med_, the emission wavelength *λ*, the quartz induced phase difference between s- and p-polarizations Δ*φ*_sp_, the OTF rescaling factors *σ*_*kx*_ and *σ*_*ky*_, the intensity ratio between the two emission paths *I*_*t*_ and the coherence strength *a*.

The refractive indices were measured from an Abbe refractometer (334610, Thermo Spectronic), which are *n*_imm_ = 1.516 for the immersion oil (16241, Cargille) and *n*_med_ = 1.351 for the SMSN imaging buffer (**Materials and Methods**).

The emission wavelength was determined by imaging a crimson bead (100 nm, custom-designed, Invitrogen), which has similar emission spectrum as the Alexa Fluor 647 (A21245, Life Technologies), at *z* positions from -1 to 1 µm with a step size of 100 nm and taking 40 frames per *z* position. An optimum *λ*, equal to 675 nm, was found by minimizing the error of the estimated step size and the stage step size, which is 100 nm. Note that this wavelength is determined by moving the sample stage, which effectively changes the thickness of the immersion medium. However, for SMSN imaging, the change of the axial position is relative to the sample medium, therefore, to account for the index mismatch, a longer wavelength of *λn*_imm_*/n*_med_ was used (**note S3**).

The phase difference Δ*φ*_sp_ *= φ*_*s*_ *-φ*_*p*_ was estimated from a stack of bead images at *z* positions from -1 to 1 µm with a step size of 10 nm. We first obtained the intensity modulations, *r*_*s*_ and *r*_*p*_, which modulate at different phases (**note S1.3**) from the bead data. We then compared those curves with the corresponding ones of the simulated PSFs with a certain Δ*φ*_sp_. By varying Δ*φ*_sp_, an optimum value, typically around -1.5 rad, was found to bring the modulation-phase difference of the simulated data close to that of the bead data.

The OTF rescaling is to account for the PSF broadening effect from isotropic dipole emission of a single fluorophore (*10, 11*). It performs as a Gaussian blur and the factors, *σ*_*kx*_ and *σ*_*ky*_ (unit: µm^−1^), are the widths of the blurring kernel in the Fourier space, which is equivalent to a Gaussian blurring kernel in real space with widths of 1/2*πσ*_*kx*_ and 1/2*πσ*_*ky*_ (unit: µm). They were estimated by minimizing the log-likelihood ratio between simulated and measured PSFs, which contains PSF images at *z* positions from -1 to 1 µm with a step size of 100 nm. Here we used 1.6 µm^−1^ for both *σ*_*kx*_ and *σ*_*ky*_. The intensity ratio and the coherence strength were discussed in detail in **note S4**.

### 3. Effect of index mismatch aberration

For oil immersion objective, the refractive index of the sample medium, e.g. 1.351 for SMSN imaging, is usually smaller than the one of the immersion oil, which is 1.516 for our 4Pi-SMSN system. This index mismatch will result in larger aberrations for imaging emitters away from the cover glass and the distortion increases with the imaging depth. Two methods can be used to incorporate the index mismatch aberration into the PR-4PiPSF model.

The first method assumes that the imaging depth is small, less than 4 µm, and within this range, the differences of PSF patterns from the ones measured at the cover glass are small (*11*). Therefore, the PSF model of emitters on the bottom cover glass was used for the entire imaging depth. To generate such a PSF model, the pupil functions, *h*_*A*_ and *h*_*B*_, were retrieved independently through the upper and lower objectives by imaging a bead on the bottom cover glass. The upper emission pupil-function *h*_*A*_ includes the index mismatch aberration caused by imaging through the entire sample medium **(fig. S10**). Therefore, the PR-4PiPSF model based on this pair of coherent pupil functions can accurately represent the emission patterns of a bead on the cover glass (**Fig. 2**). However, in SMSN imaging, the defocused PSFs were formed by emitters at different axial locations relative to the sample medium instead of by moving the sample stage, which changes the thickness of the immersion medium in the case of bead imaging. The modulation period of the emission patterns in the sample medium is larger than the one in the immersion medium. Therefore, a factor of *n*_imm_*/n*_med_ is multiplied to the estimated emission wavelength from bead imaging (**note S2**).

The second method is to build a PSF model that can model the emission patterns came from arbitrary axial locations inside the sample medium. To achieve this, we first excluded index mismatch aberration from the phase retrieved pupil functions: the upper emission pupil-function *h*_*A*_ was retrieved from imaging a bead on the top cover glass through the upper objective, while the lower emission pupil-function *h*_*B*_ was retrieved from imaging a bead on the bottom cover glass through the lower objective (**fig. S10**). Next, we derived a PSF model incorporating index mismatch aberration by conducting a ‘gedanken-experiment’ as the following: first, let both objectives focus on the cover glass close to themselves and then move the upper objective a distance of *H* to focus on the bottom cover glass; second move the sample stage a distance of *d* to focus on the target image plane inside the specimen, then the upper and lower emission pupil functions describing emitters at this image plane can be modeled as,

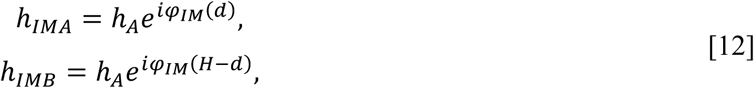

where *φ*_*IM*_ is the aberration phase caused by index mismatch (*11*),

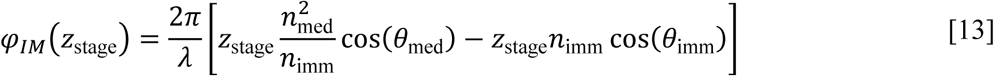

and it varies with the stage position, *z*_stage_, which is the distance of the stage moved to focus on a plane away from the designed focal plane of the objective. Note that the value of *z*_stage_ is relative to the immersion medium, in the case of 4Pi-SMSN system, there are two stage positions, *d* and *H − d*, where *H* is a constant during imaging and those two quantities were only varied by the distance *d* the sample stage moved. Therefore, at a particular *d* the index mismatch aberrations *φ*_*IM*_ are fixed, the emission patterns are only dependent on the emitters’ axial locations *z* inside the sample medium,

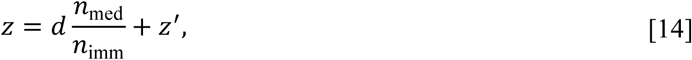

which is zero at the bottom cover glass, and *z*^*′*^ is a relative position (*11*) to the image plane position *dn*_med_*/n*_imm_. Together by replacing *h*_*A*_ and *h*_*B*_ with *h*_*IMA*_ and *h*_*IMB*_, and change the defocus term to

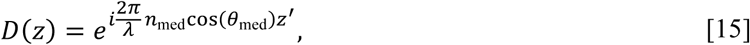

in **Main Text** Eq. 1, the 4Pi-PSF model at arbitrary axial positions can be calculated. The wavelength *λ* used here is the one measured from bead imaging, which is approximate to the emission wavelength in air (**note S2**).

We found that the PSF models generated from both methods are consistent in a large axial range (**fig. S13**). In this work, we used the first method for PSF modeling because all the presented data were acquired within a small imaging depth. For thick sample imaging, the second method could be used to account for the depth-dependent index mismatch aberration.

### 4. Effect of intensity ratio and coherence strength

As described in the PR-4PiPSF generation (**Main Text**), both the intensity ratio *I*_*t*_ and the coherence strength *a* can affect the modulation depth of the 4Pi-PSF and consequently influence the localization precision. We investigated the theoretical localization precision of 4Pi-PSFs with various coherence strength and intensity ratios based on CRLB (with 5000 total photon and 5 background photon, **fig. S11**): 1) When intensity ratio is zero, the system reduces into a single-objective microscope where the localization precisions are the worst; 2) When keeping the coherence strength at zero but increasing the intensity ratio, the precision can be ideally improved by a factor of 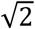 predicting the performance of an incoherent dual-objective microscope (*12*); 3) Only when increasing both the intensity ratio and the coherence strength, the precision can be further improved. We found that the axial precision is more sensitive to the coherence strength and can be improved by a factor of 20 comparing with single objective cases. We also noticed that, the increasing speed of the axial precision slows down when the intensity ratio and coherence strength are sufficiently large, such as within the dark blue region bounded by an axial precision of 4 nm. This indicates that 4Pi-PSFs, in terms of precision, have adequate tolerance with these two factors. For example, with high intensity ratio, a moderate drop in coherence strength will not drastically deteriorate the resolution, which is especially important for thick sample imaging, where the coherence strength is depth dependent. In this work, we empirically set intensity ratio to 1 and coherence strength to 0.8.

### 5. Temperature modulation on 4Pi-PSFs

To quantify the effect of temperature fluctuation on 4Pi-SMSN system, we measured the interference phase *φ* from bead imaging (**note S1.3**) and simultaneously collected the current temperature using a temperature sensor (TSP01, Thorlabs) at a sequence of 500 time points with 10 s intervals. From the measured interference phase *φ* and together with the axial position *z* of the emitter, localized with astigmatism method, the effective axial drift caused by the change of cavity phase *φ*_0_ can be calculated from,

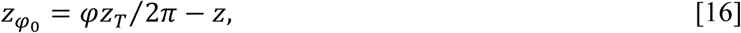

where *z*_*T*_ is the modulation period of the 4Pi-PSF with respect to the immersion medium and the cavity phase *φ*_0_ can be estimated from 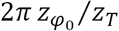. We found that the fluctuation of 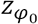 closely follows the temperature fluctuation, which indicates that the cavity temperature has noticeable effect on the emission pattern of 4Pi-SMSN systems (**fig. S5**).

### 6. Quadrant alignment

Data for quadrant alignment are images of sparse beads sample with 5-10 beads per field of view (FOV) or cell sample illuminated with a low laser power (**Materials and Methods**). Each of those datasets contains 40-100 frames collected from the lower emission path. A previously developed alignment procedure (*1*) was applied: 1) Four quadrants were cropped from the raw camera frames and each measures 168 × 168 pixels; 2) Select one quadrant as the reference, and find the linear transformations, including scaling, translation and rotation, from each quadrant to the reference quadrant using the Dipimage (*13*) functions *fmmatch* and *find_affine_trans* sequentially; 3) Align each quadrant to the reference quadrant using the Dipimage function *affine_trans* given the obtained transformation relations.

For PR-4Pi method, to achieve maximum likelihood estimation (MLE), the statistical behavior (mean and variance) of the pixel value should remain unchanged. Therefore, instead of transforming the raw data, we directly incorporated the transformation relationships between quadrants inside the likelihood function during MLE. Specifically, during the segmentation procedure (**fig. S7**), we generated two sets of subregions, one was from aligned quadrants and used for cavity phase calibration and initial guess for the axial positions, and another one was from unaligned quadrants and used for MLE. The segmentation procedure for unaligned quadrants was as follows: 1) Select candidate emitters from the sum of the aligned quadrants and record the pixel coordinates of the candidate emitters as *coords1*. 2) Transform *coords1* to quadrants 2 to 4, assume quadrant 1 is the reference, and record the results as *coords2, coords3* and *coords4*. 3) Crop subregions in each unaligned quadrant with the nearest integer of the corresponding *coords* as the centers. The variance and gain maps for each subregion were also cropped during this step. 4) Record the local shift of each subregion as

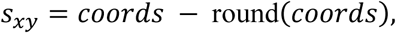

which will be applied to the likelihood function during MLE.

### 7. Linear and cubic interpolations in PR-4Pi algorithm

The speed of PR-4Pi localization method depends majorly on the speed of generating PR-4PiPSF models, each requiring multiple Fourier transforms. Therefore, we replaced the Fourier transform with interpolation and implemented it with MLE (**note S1.3** and **fig. S7**) on GPU to speed up the regression process. The interpolation algorithm generates 4Pi-PSFs at arbitrary locations by interpolating a pre-generated PR-4PiPSF library, PSFlib, which covers an axial range of -1.5 to 1.5 µm and a lateral extend of 25 × 25 camera pixels, and each voxel of the PSFlib measures 0.25 pixel × 0.25 pixel × 5 nm. Together with the generation of the 4Pi-PSF model, whose first and second derivatives were also calculated during the interpolation process.

We have implemented both linear and cubic interpolation methods in the PR-4Pi algorithm and we found that linear interpolation tends to produce larger artifact when operating on complex PSF patterns such as 4Pi-PSFs (**fig. S14**). However, both methods achieve comparable resolutions on localizing either bead or SMSN data (not shown). The GPU implementations of linear and cubic interpolations achieve a localization speed of 1140 emitter/s and 270 emitters/s respectively, nearly 600 times faster than the corresponding CPU implementations using MATLAB (MathWorks, Natick, MA). In this work, cubic interpolation was used for all the demonstrations on PR-4Pi algorithm.

#### 7.1 Justification on using 100 nm bead

Here we used 100 nm bead for phase retrieval for its high photon count. We found a few literatures using 100 nm for PSF measurements. It is true that 100 nm is relatively large comparing with the size of a single molecule, however, it is still well below the diffraction limit. A large bead size will have a broadening effect on measured PSFs and therefore could shrink the retrieved pupil size, however, our retrieved pupil fills the entire aperture defined by the objective NA (**Fig. 1** and **fig. S13**), which indicates that 100 nm bead is capable of retrieving features of the entire pupil. Furthermore, dipole radiation of a single molecule will also broaden the PSFs, so that we have introduced a OTF rescaling factor to account for this effect (*10, 11*) (**note S3**). However, we found that for localizing bead data with spline interpolation, the OTF rescaling factor has little effect on the localization results (**fig. S15**). Only when σ_OTF_ (*σ*_*kx*_ and *σ*_*ky*_) equal to 11 µm^−1^, corresponding to a smooth kernel of 0.1 pixels (pixel size at 129 nm) in real space (almost no smoothing), the localization results start to show divergence at a few positions. Therefore, in our opinion, using 100 nm bead is justifiable for phase retrieval as it doesn’t sacrifice the pupil size and its PSF broadening effect has ignorable variations on localization results in practice.

#### 7.2 Linear interpolation

Consider a one-dimensional (1D) situation, a linear interpolation between sample positions *x*_1_ and *x*_2_ is,

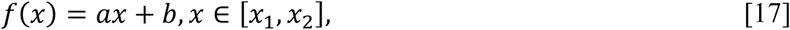

where *a* and *b* are interpolation coefficients. Given the function values, *f*_1_ and *f*_2_, at the sample positions, solve the following linear equations for *a* and *b*,

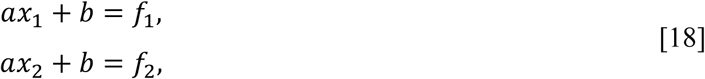

we have

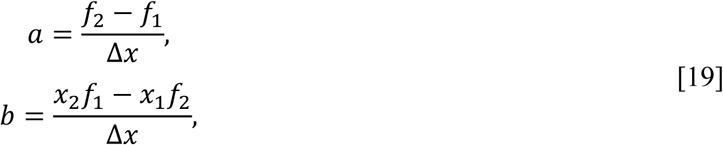

where Δ*x* is the distance between the sample positions and equal to *x*_2_ − *x*_1_. The first and second derivatives of the interpolation function *f*(*x*) is *a* and 0 respectively.

To generalize above calculations to three dimensions, we used a chain of 1D interpolations to obtain the function value at any location inside a voxel defined by the sample positions (*x*_1_, *y*_1_, *z*_1_) and (*x*_2_, *y*_2_, *z*_2_). The 1D interpolation functions were calculated in the order of *x* dimension, followed by *y* and *z* dimensions,

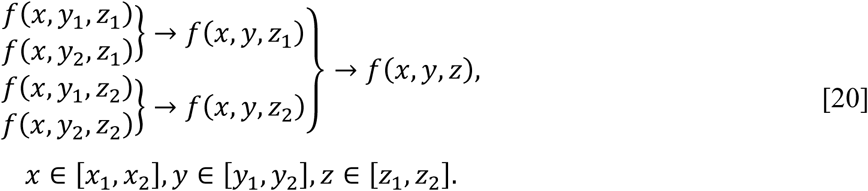

The interpolation functions at each dimension can be written as,

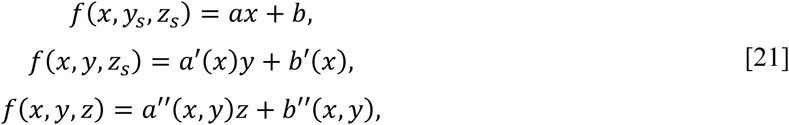

where *s* denotes the sample positions. Therefore, the first derivative of *f*(*x, y, z*) with respect to *x, y* and *z* can be also calculated from a chain of 1D interpolations in the same order,

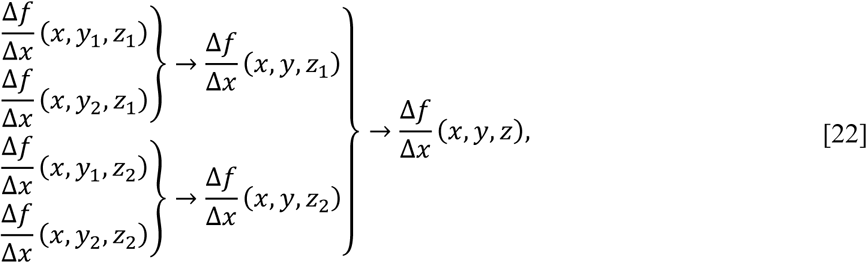

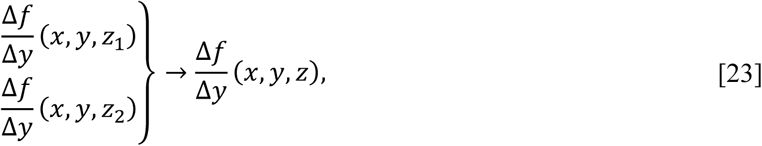

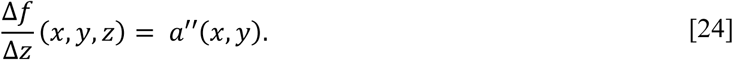

The second derivatives are zero. By evaluating a series of 1D linear interpolations, we can obtain the function value at any location inside a given voxel as well as its first and second derivatives with respect to each dimension.

#### 7.3 Cubic interpolation

Consider a one-dimensional (1D) situation, a cubic interpolation between sample positions *x*_1_ and *x*_2_ is,

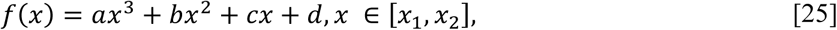

where *a, b, c* and *d* are interpolation coefficients. To obtain those four coefficients, two more sample points are required, one preceding *x*_1_ and one after *x*_2_, which are *x*_0_ and *x*_3_. Given the function values at the four sample positions, *f*_0_, *f*_1_, *f*_2_ and *f*_3_, solve the following linear equations for *a, b, c* and *d*,

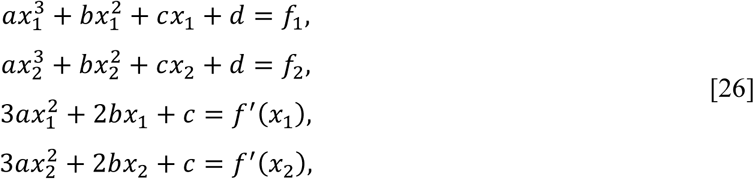

we have

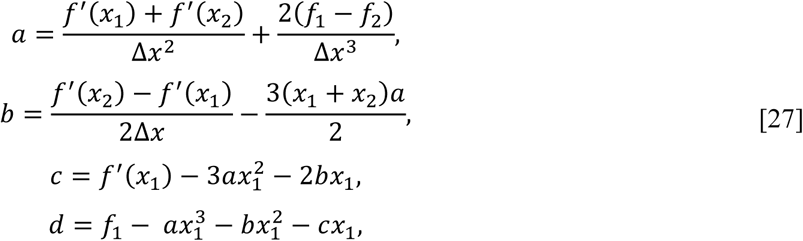

where Δ*x* is the distance between adjacent sample points, and the first derivatives of *f (x)* at *x*_1_ and *x*_2_ were calculated from the central discrete derivative,

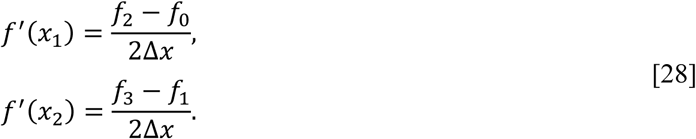

Then the first and second derivatives of the interpolation function are,

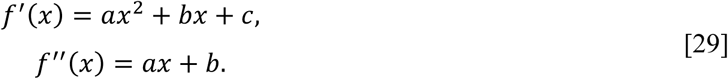

To generalize above calculations to three dimensions, we used a chain of 1D interpolations to obtain the function value at any location inside a voxel defined by the sample positions (*x*_1_, *y*_1_, *z*_1_) and (*x*_2_, *y*_2_, *z*_2_). The 1D interpolation functions were calculated in the order of *x* dimension, followed by *y* and *z* dimensions,

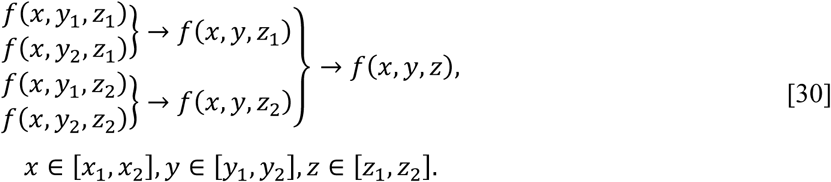

The interpolation function at each dimension can be written as,

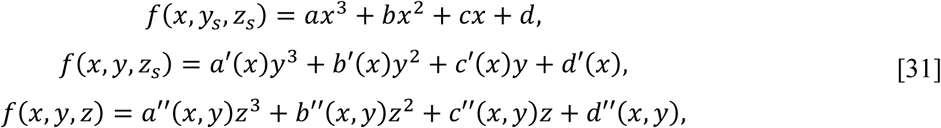

where *s* denotes the sample positions. Therefore, the first derivative of *f(x, y, z)* with respect to *x, y* and *z* can be also calculated from a chain of 1D cubic interpolations in the same order,

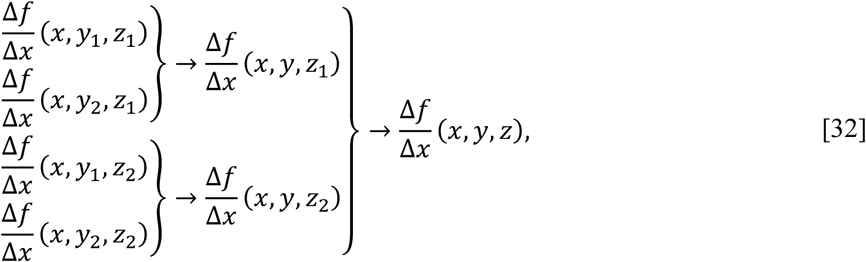

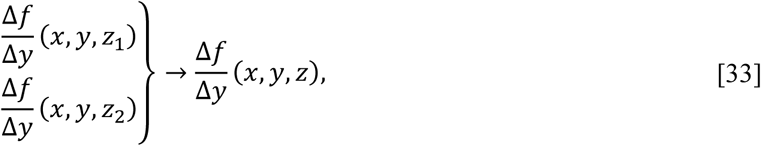

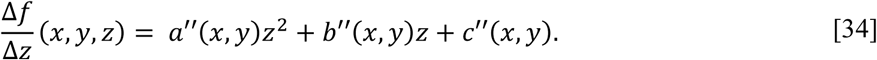

Similarly, we can obtain the second derivatives of *f(x, y, z)*. By evaluating a series of 1D cubic interpolations, we can obtain the function value at any location inside a given voxel as well as its first and second derivatives with respect to each dimension. With this method, there is no need to store a huge number of interpolation coefficients (*14, 15*), 64 for each voxel, to generate PSFs at arbitrary locations. And the evaluation of a 1D cubic interpolation is sufficiently fast.

### 8. Increase of information content using interferometric PSF

Here we provide a generic proof on the increase of the information content by using an interferometric PSF. Consider a simple interferometric system that has two channels, the formation of its PSF can be considered as a redistribution of the emitted photons of a single emitter into two channels. The sum of the PSF patterns from the two channels produces a conventional incoherent PSF,

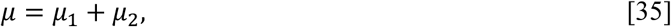

where the PSF of each channel is

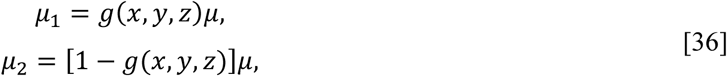

where *g(x, y, z)*, the same size of *µ*, is a function of the emitter’s *x, y* and *z* positions with pixel values at 0 to 1. This function can be considered as a redistribution function of the emitted photons. To quantify the information content of a PSF pattern, we use the Fisher information matrix (*16*). Under the assumption that the pixel values follow a Poisson process and are independent, the Fisher information matrix of an incoherent PSF is (*16*)

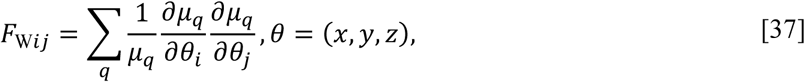

where *q* is the pixel index. And the Fisher information matrix of a two-channel interferometric PSF is (*11*)

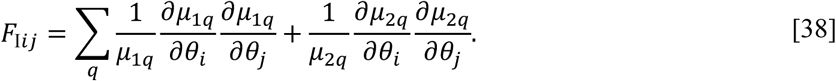

Considering only the diagonal terms of the Fisher information matrix, which are the dominant factors when all estimation parameters in *θ* are independent, the Fisher information of pixel *q* in *F*_I_ is

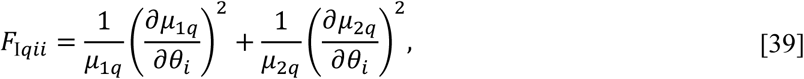

substitute Eq. **36** to above equation, we have,

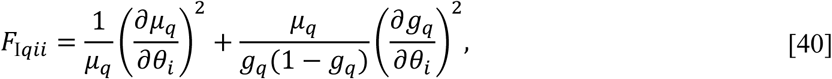

where the first term represents the information of the incoherent PSF and the second term, always greater than zero, is the extra information from the interferometric PSF. From this expression, the larger the gradient of the redistribution function *g(x, y, z)* along a certain dimension, the higher the increase of the information associated with that dimension.

### 9. Quantification of light sheet profile

To quantify the thickness of the light sheet, a sample with fluorescent dye (A20006, Life Technologies) coated coverslip was prepared (**Materials and Methods**). The light sheet direction was adjusted to be along the axial direction. Subsequently, using a piezo nano-positioner sample stage (P-541.ZCD, Physik Instrumente, Germany), the thin layer of dye was moved at various axial positions and imaged through the upper objective in focus. This method utilizes the special geometry of 4Pi systems where the upper objective can be moved along the axial direction independently without changing the shape of the light sheet which was generated from the lower objective. The fluorophore image at each axial position was averaged along the laser line direction (the illumination profile is similar as a wide laser line) and the resulting intensity profile was fitted with a Gaussian function. The width of the fitted Gaussian at its amplitude of 1/e is the width of the light sheet at the measured axial position. Then the thickness of the light sheet, the smallest width (2*ω*_0_, where *ω*_0_ is the beam waist of a Gaussian beam (*17*)), was estimated by fitting the obtained widths at various axial positions with a Gaussian beam profile (*17*). Importantly, when the light sheet is tilted, because of the refractive index (*n*) mismatch between the sample medium (e.g. *n* is 1.351) and immersion oil (e.g. *n* is 1.516), the sheet thickness changes, which can be estimated as 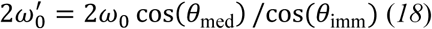 (*18*), where *θ*_imm_ is the incident angle in the immersion oil and *θ*_med_ is the refraction angle in the sample medium. Therefore, the larger the tilted angle the thinner the inclined light-sheet, we found that a ∼2.3 µm light-sheet thickness after inclining 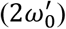 can be achieved with *θ*_med_ at 65° and *2ω*_0_ at 3.2 µm (**fig. S8**)

The above obtained light-sheet thickness is along the light propagation direction, from the imaging point of view, the confinement of an inclined light-sheet can be characterized by its axial extent relative to the sample plane, which can be approximated to 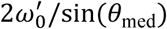. The illumination area of the inclined light-sheet, defining the field of view (FOV), is a rectangular region with one dimension the same size as for the epi illumination and the other dimension approximating to 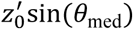, where 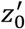 equal to 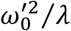 is the confocal parameter (*17*) of a Gaussian beam with a beam waist of 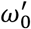 and a wavelength of *λ*. The estimated FOV of the light-sheet illumination in our 4Pi-SMSN system is 22 × 8 µm^2^ given the excitation wavelength (*λ*) at 642 nm.

## Supplementary Figures

**Fig S1.**
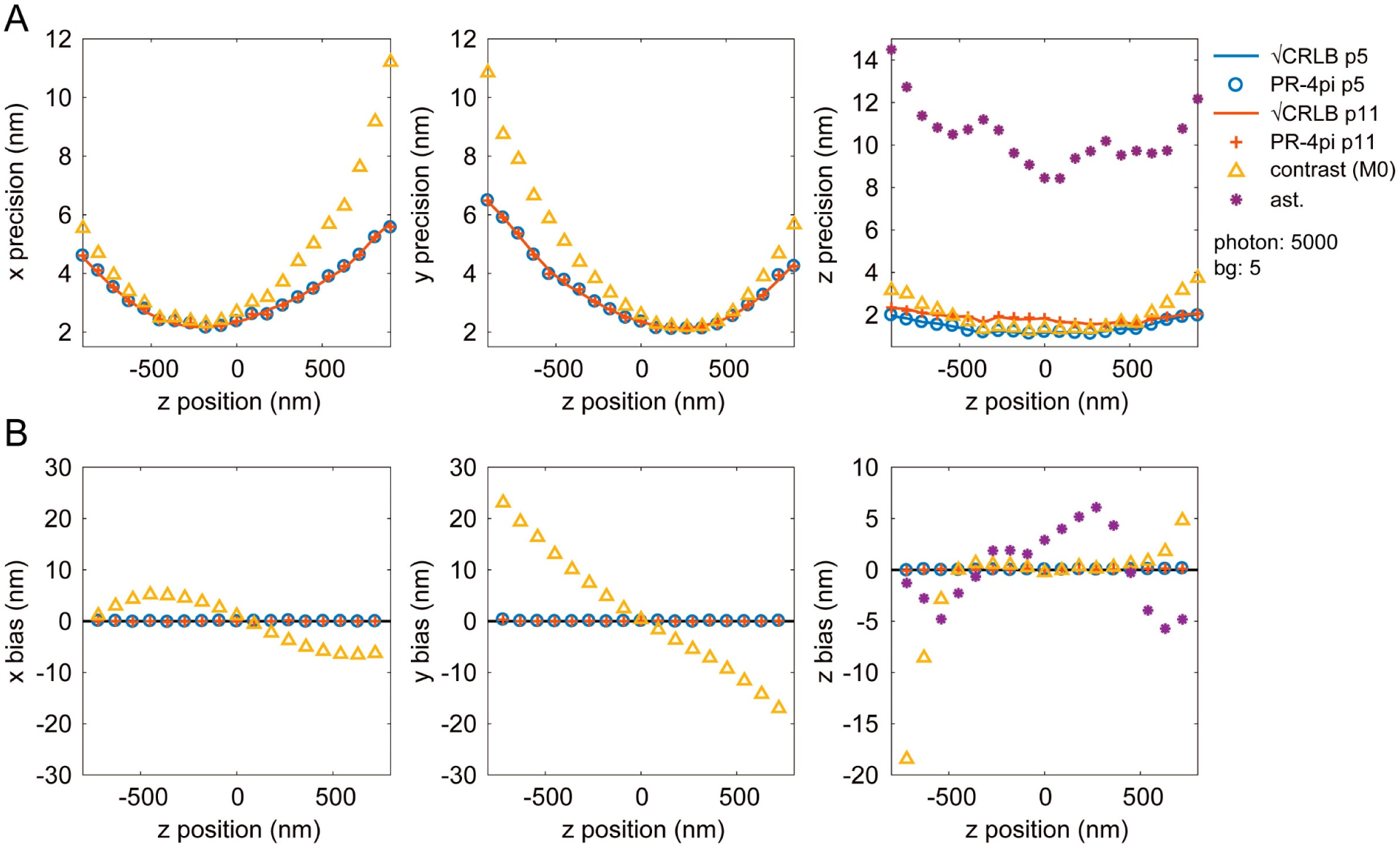
Comparison of various localization methods with CRLB on simulated 4Pi-PSF data with astigmatism modification. PR-4Pi algorithms with fixed (PR-4Pi p5, 5 parameters) and independent (PR-4Pi p11, 11 parameters) intensity and background between four quadrants were compared. Contrast method refers to the algorithm developed in (*1*) that extracts the axial positions using the Gaussian-weighted 0^th^ central moment (M0, **note S1.3**). The astigmatism method (ast.) is based on a Gaussian PSF model and operated on incoherent PSFs that are the sum of the 4Pi-PSFs of the four channels (**Fig. 1**). Data were simulated from phase retrieved pupil functions with enhanced astigmatism aberration and contain 1000 PSFs at each axial position in the range of -900 nm to 900 nm, with a step size of 100 nm. The total photon and background photon used were 5000 per objective and 10 per pixel. Note that, for **Fig. 4** and **figs. S1-S4**, in order to generate localization results that are information limited without the artifact caused by ghost imaging, initial guesses that are close to global minimums were used for PR-4Pi algorithms. And for methods using Gaussian-weighted central moments, since the actual axial position increases monotonically with the frame number, the obtained phases can be easily unwrapped with MATLAB function *unwrap* and therefore eliminated ghost images, which however is not possible for cell data. Here we used ‘bias’ for simulated data as the ground truth is known and ‘deviation’ for bead data. In all figures, without specification, PR-4Pi denotes the algorithm using 11 estimation parameters.

**Fig S2.**
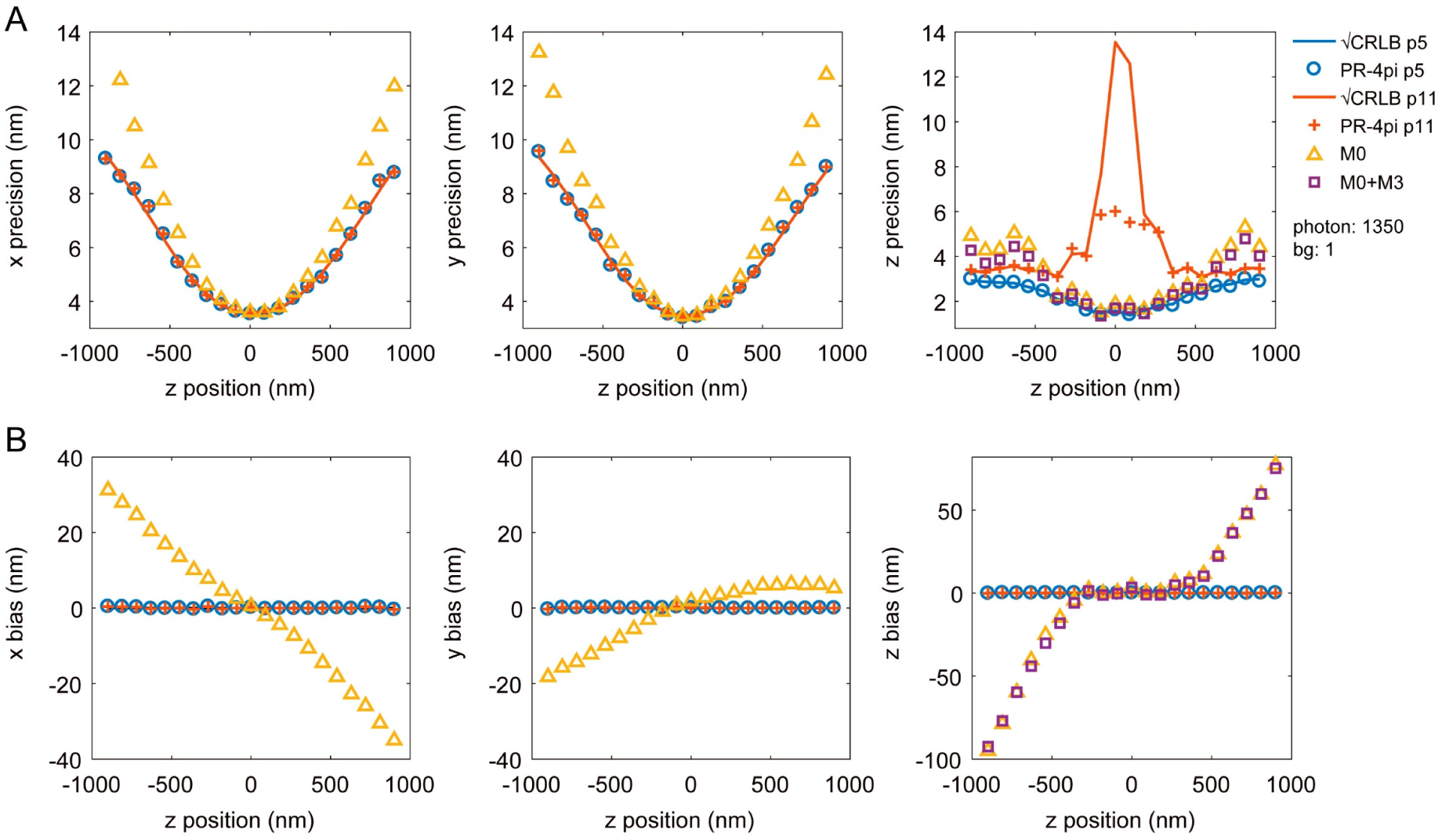
Comparison of various localization methods with CRLB on simulated 4Pi-PSF data *without* astigmatism modification. PR-4pi algorithms with fixed (PR-4Pi p5) and independent (PR-4Pi p11) intensity and background between four quadrants were compared. The axial positions were also extracted with Gaussian-weighted central moments, where M0 used the 0^th^ central moment and M0+M3 used a combination of 0^th^ and 3^rd^ central moments as described in (*5*). We also showed the lateral localization results (orange triangles) using a Gaussian PSF model on incoherent PSFs that are the sum of the 4Pi-PSFs of the four channels (**Fig. 1**). Data were simulated from phase retrieved pupil functions without astigmatism modification and contain 1000 PSFs at each axial position in the range of -900 nm to 900 nm, with a step size of 100 nm. The total photon and background photon used were 1350 per objective and 1 per pixel. Note that here we simulated the PSF data without astigmatism and with a photon count comparable with the bead data demonstrated in (*5*) (**table S1**), in order to have a fair comparison with precious methods.

**Fig S3.**
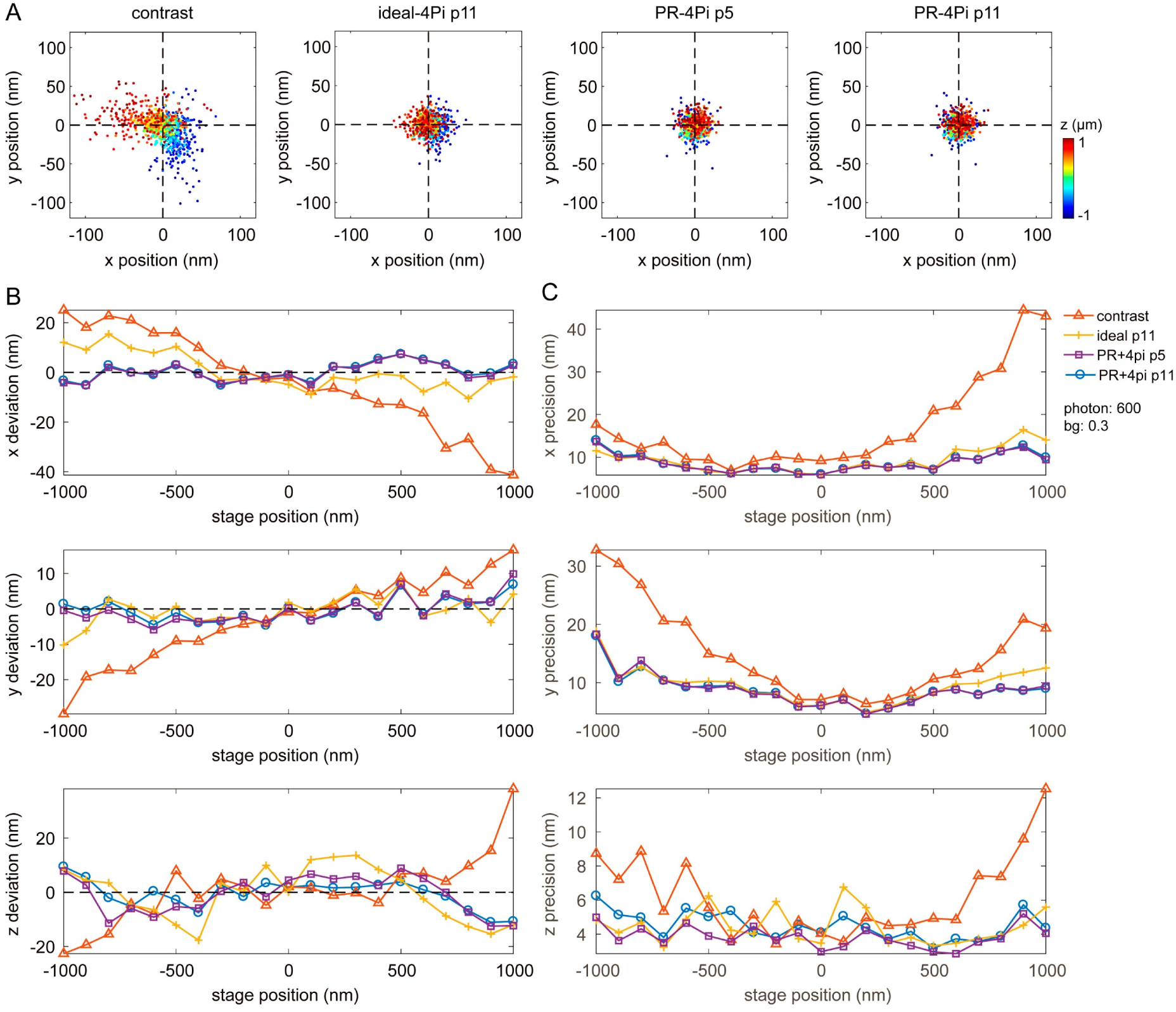
Comparison of localization results using various PSF models on experimental bead data with low photon count. (**A**) Scatter plot of lateral localizations from an axial scan of 100 nm bead from 1 µm above to 1 µm below the common focus of two 4Pi objectives. Estimations with fixed (p5, 5 parameters) and independent (p11, 11 parameters) intensity/background between four quadrants were compared. Color scaled with axial positions. (**B**) Localization deviations in *x, y* and *z*. (**C**) Localization precisions in *x, y* and *z*. Comparing with PR-4Pi p11, PR-4Pi p5 improves the axial precisions but sacrifices the accuracy (with increased axial deviations). Data were acquired by imaging a fluorescence bead at axial positions from -1 to 1 µm by translating the sample stage with a step size of 100 nm, 40 frames were captured at each axial position (**Fig. 4** and **Materials and Methods**). The PR-4PiPSF models were generated by phase retrieved pupil function and the ideal-4PiPSF models used unaberrated pupil functions and assumed no transmission loss and beam splitting inequality presented in the imaging system. The estimation results from PR-4Pi method yielded a mean total photon of 600 per objective and a background photon of 0.3 per pixel. Note that **figs. S3** and **S4** used the same PSF models for PR-4Pi and ideal-4Pi, and the bead data used for phase retrieval are always different from the bead data used for localization.

**Fig S4.**
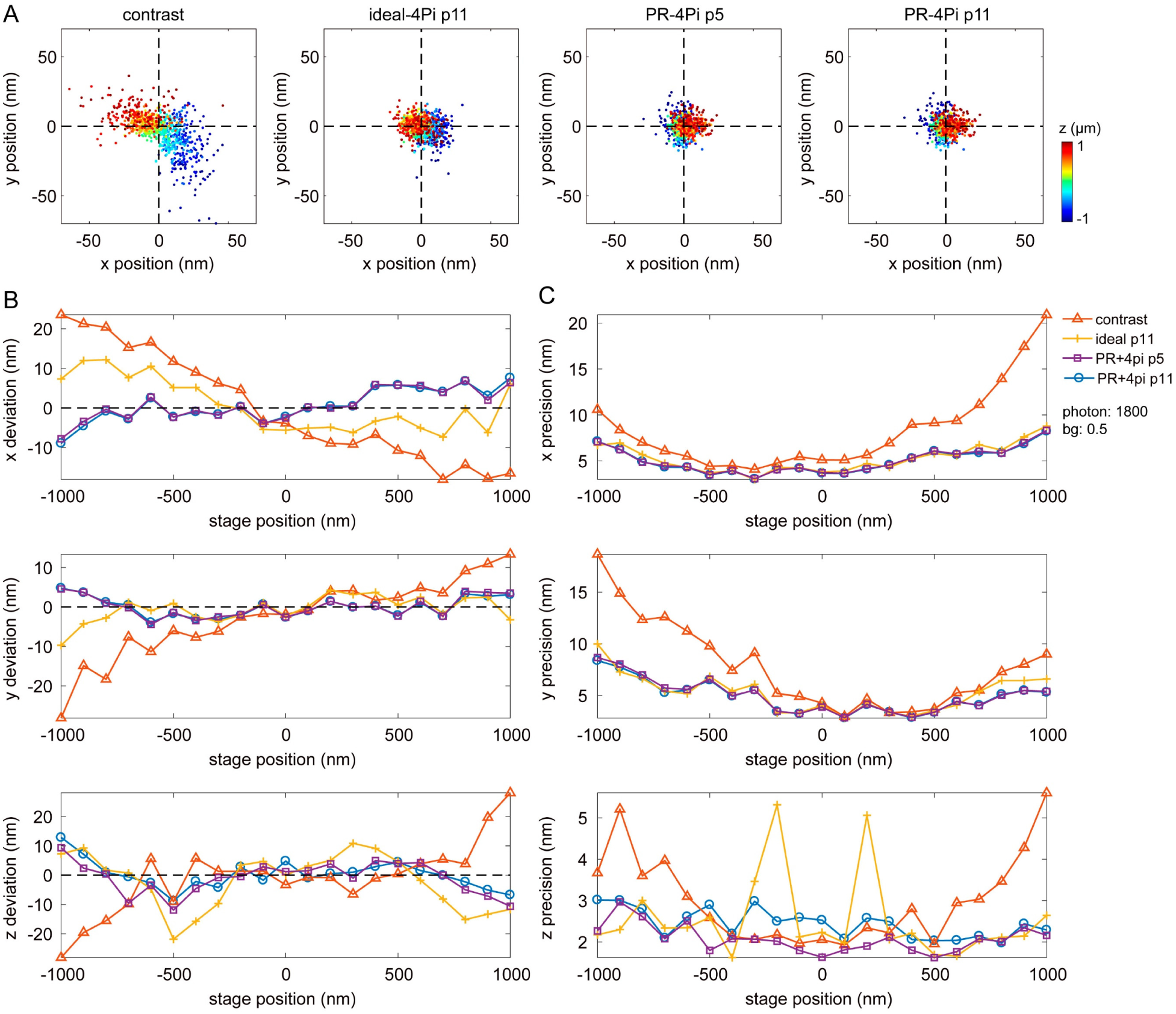
Comparison of localization results using various PSF models on experimental bead data with mid-level photon count. (**A**) Scatter plot of lateral localizations from an axial scan of 100 nm bead from 1 µm above to 1 µm below the common focus of two 4Pi objectives. Estimations with fixed (p5, 5 parameters) and independent (p11, 11 parameters) intensity/background between four quadrants were compared. Color scaled with axial positions. (**B**) Localization deviations in *x, y* and *z*. (**C**) Localization precisions in *x, y* and *z*. Comparing with PR-4Pi p11, PR-4Pi p5 improves the axial precisions but sacrifices the accuracy (with increased axial deviations). Data were acquired by imaging a fluorescence bead at axial positions from -1 to 1 µm by translating the sample stage with a step size of 100 nm, 40 frames were captured at each axial position (**Fig. 4** and **Materials and Methods**). The PR-4PiPSF models were generated by phase retrieved pupil function and the ideal-4PiPSF models used unaberrated pupil functions and assumed no transmission loss and beam splitting inequality presented in the imaging system. The estimation results from PR-4Pi method yielded a mean total photon of 1800 per objective and a background photon of 0.5 per pixel.

**Fig S5.**
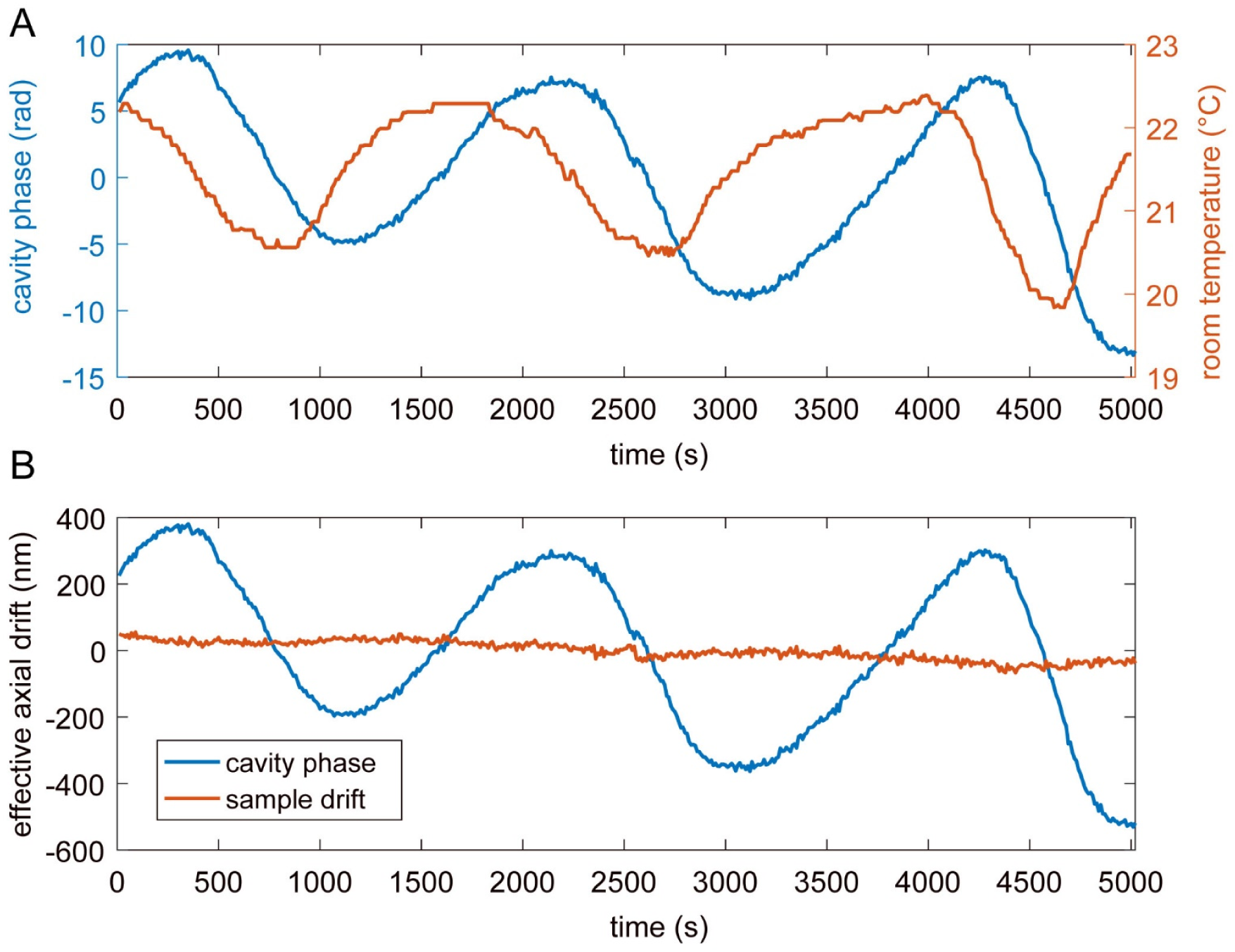
Correlation of temperature fluctuation and cavity phase drift. The room temperature was measured using a temperature sensor (TSP01, Thorlabs, resolution at 0.05 °C) at a sequence of 500 time points with 10 s intervals, and at the same time the cavity phase drift was measured by imaging a fluorescence bead (**note S5**). (**A**) Fluctuations of cavity phase and room temperature show strong correlations. (**B**) The effective axial-drift measurements from cavity phase change and the sample drift. The cavity-phase induced axial-drift is the main factor that deteriorates the axial resolution of 4Pi-SMSN systems.

**Fig S6.**
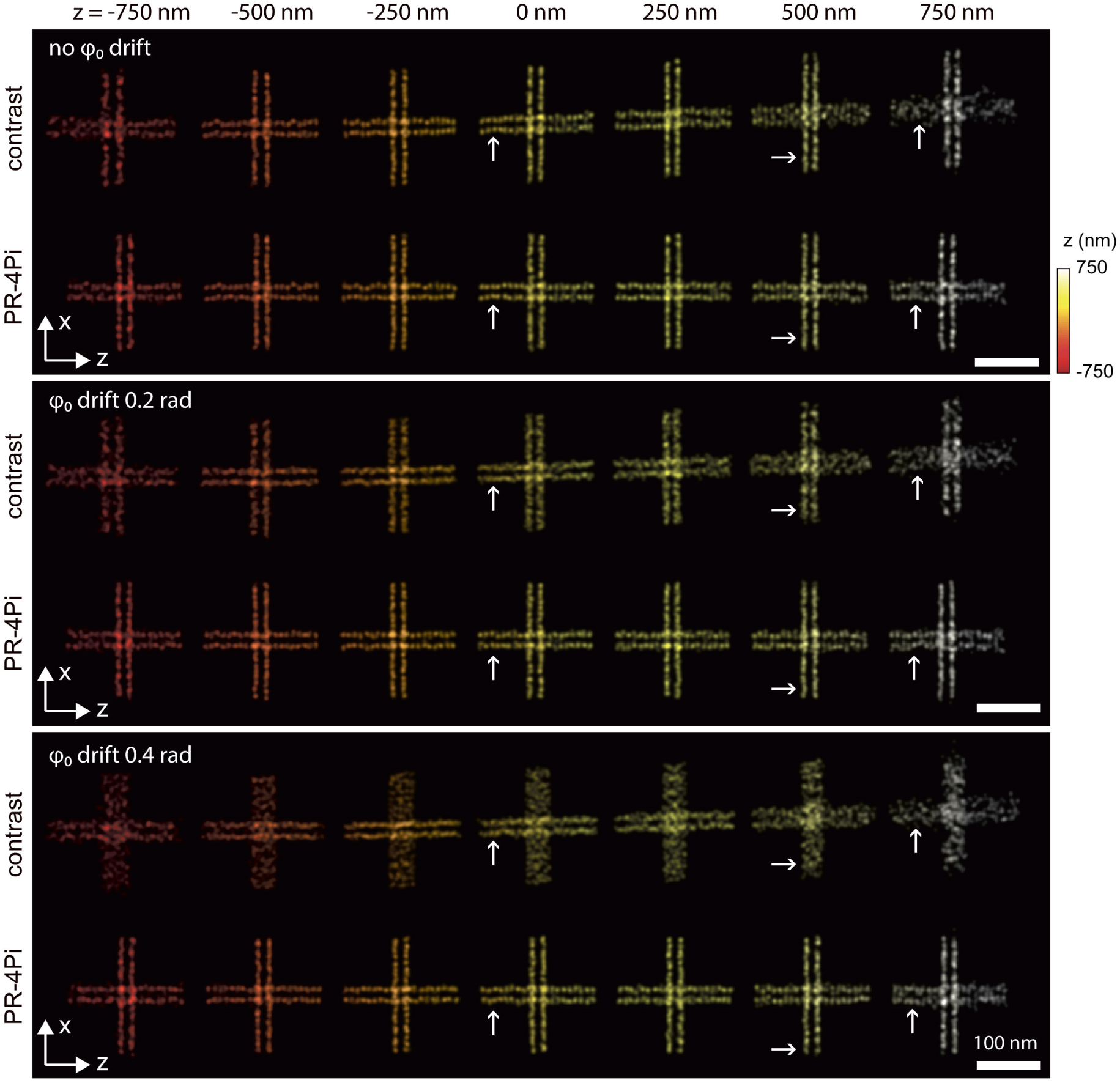
Effect of cavity phase drift on axial resolution. A double-line cross structure at the x-z plane was simulated at various axial positions. Each line measures 200 nm in length and contains 20 points spaced uniformly, and the double lines are separated by 20 nm. Five PSFs with cavity phase (*φ*_0_) equally spaced between 0 up to 0.4 rad were generated at each point of the simulated structures using phase retrieved pupil functions. We found that in most cases, PR-4Pi succeeded in precise localization of these 20 emitters per line resulting in almost equally spaced and distinct dots along the structure. In comparison, contrast based method manages to resolve these ultra-structures when it is close to the common focus with 0 cavity phase drift (first row, *z* at 0 nm), but fails to resolve these ultra-structures in most cases. In lateral dimensions, the contrast based method results in larger deviations and worse resolutions at out-of-focus positions. In the axial dimension, the contrast based method fails to resolve the double lines under a *φ*_0_ drift of 0.4 rad, which is below the average cavity phase drift (0.45 rad, measured from our system) within 40 s, the shortest time segment for post drift correction. Data were simulated from phase retrieved pupil functions, and the total photon and background photon used were 5000 per objective and 10 per pixel.

**Fig S7.**
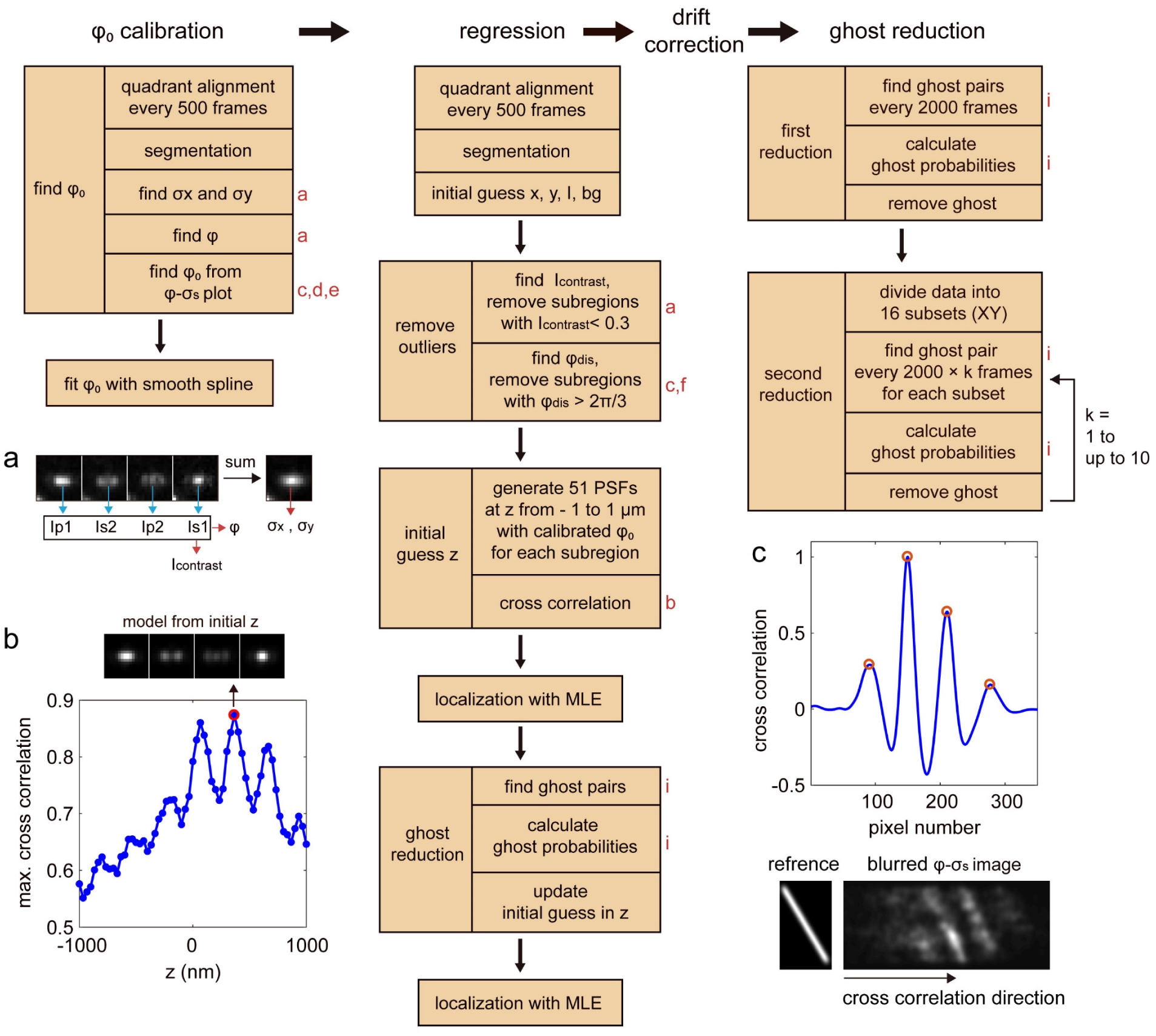
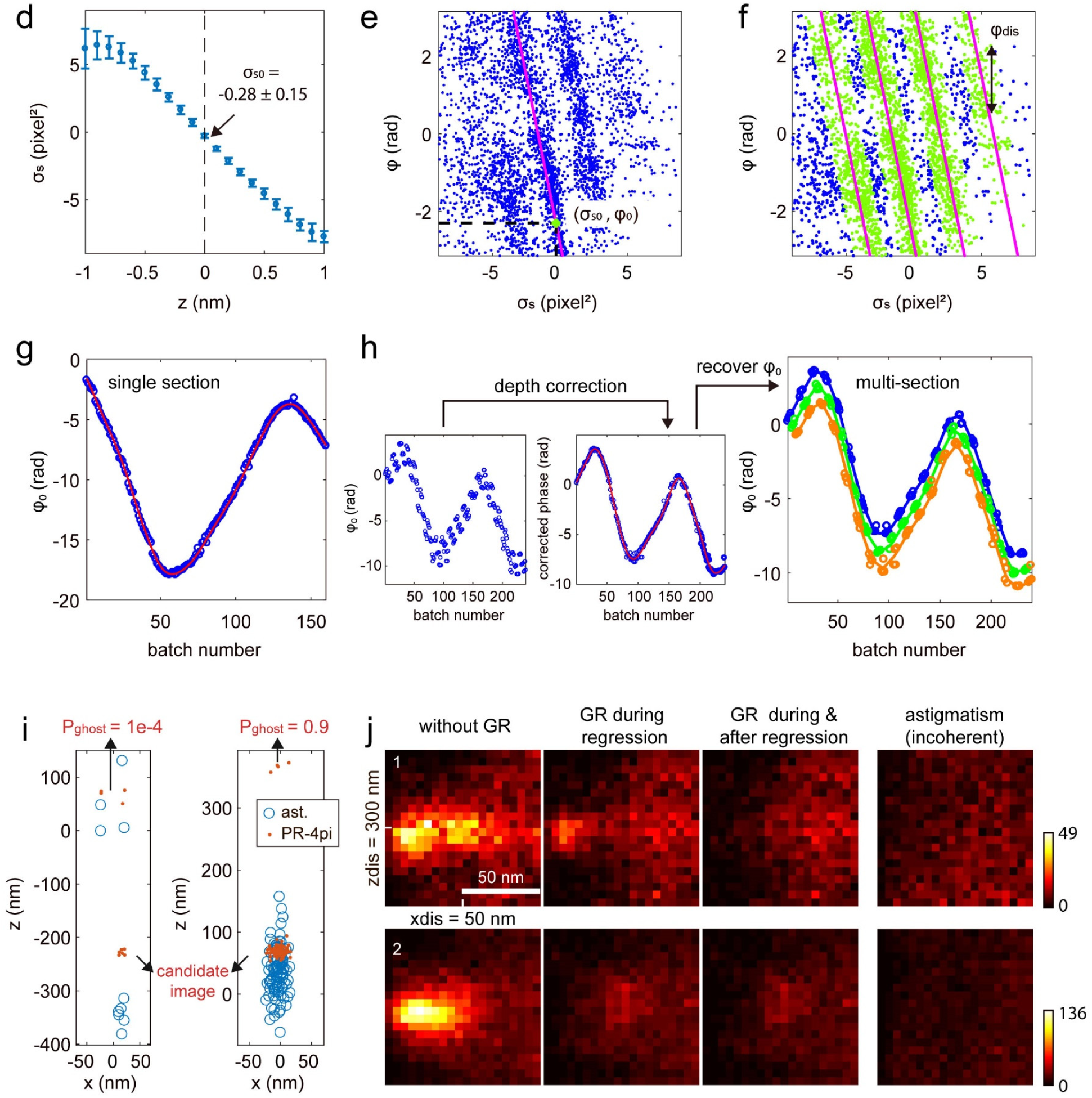
Flow chart of PR-4Pi localization algorithm on SMSN data. Steps marked with red letters on the right are accompanied with corresponding graphic illustrations following the flow chart. (**a**) An example of 4Pi-PSF-image of a single emitter, it contains PSFs from four channels (P1, S2, P2 and S1), from which the interference phase (*φ*) and the modulation contrast (*I*_contrast_) can be estimated (**note S1.3**). And the sum of the PSFs from the four channels was used to estimate the PSF widths *σ*_*x*_ and *σ*_*y*_ (**note S1.3**). (**b**) Cross correlation of a 4Pi bead-image with a set of library 4Pi-PSFs. At each axial position, first the maximum cross correlation value between the bead-image and the library 4Pi-PSF from each of the four channels was calculated, then the mean of the obtained four values was used to generate the curve. The optimum *z* position (red circle) was used as the initial z position for maximum likelihood estimation (MLE). (**c**) Cross correlation of a reference line-image and a Gaussian blurred *φ*-*σ*_*s*_ image along the *σ*_*s*_ dimension. Peaks above a set threshold were selected (red circles) to determine the fitted line positions in the *φ*-*σ*_*s*_ plot (e,f). **Fig. S7 continued.** (**d**) The shape metrics of the 4Pi-PSFs measured at *z* positions from -1 to 1 µm, with 100 nm intervals. The error bar at each axial position presents the standard deviation of the measurements from five bead data. (**e**) Estimation of the cavity phase, *φ*_0_. It was estimated from the line fit (magenta line) of the stripe with the highest emitter density, selected from the highest peak of the cross correlation result in (c). (**f**) Removal of outliers (blue dots) that have a *φ*_dis_ greater than 2π/3, which was measured by the vertical distance of each point to its nearest line fit (magenta lines). (**g**) Cavity phase calibration from single-plane imaging of mitochondria (TOM 20) in COS-7 cell. Cavity phase *φ*_0_ for every data batch (10 s each) were extracted and subsequently its time evolution was obtained through interpolation (**note S1.3**). (**h**) Cavity phase calibration from multi-plane imaging of mitochondria (TOM 20) in COS-7 cell. The initial estimates of *φ*_0_ were depth corrected to form a smooth phase-evolution and fitted with a smooth spline function. The actual evolution of *φ*_0_ for each plane was then recovered from the fitted spline. (**i**) Examples of ghost pairs after regression. Not all ghost pairs are from the same emitter, only the ghost images with high ghost probabilities (P_ghost_) were updated or removed (**note S1.4**). (**j**) Density of ghost pairs with or without ghost reduction (GR), where zdis and xdis represent the pairwise distances of localized positions in axial and lateral dimensions respectively (**note S1.4**). Ghost images were largely reduced by GR during the regression step and further diminished by GR after regression. The resulting ghost density is comparable with the one under incoherent detection, which is ghost free. Incoherent PSFs were generated from the sum of the 4Pi-PSFs of the four channels and were localized with astigmatism method (ast.) based on a Gaussian PSF model. (**note S1**).

**Fig S8.**
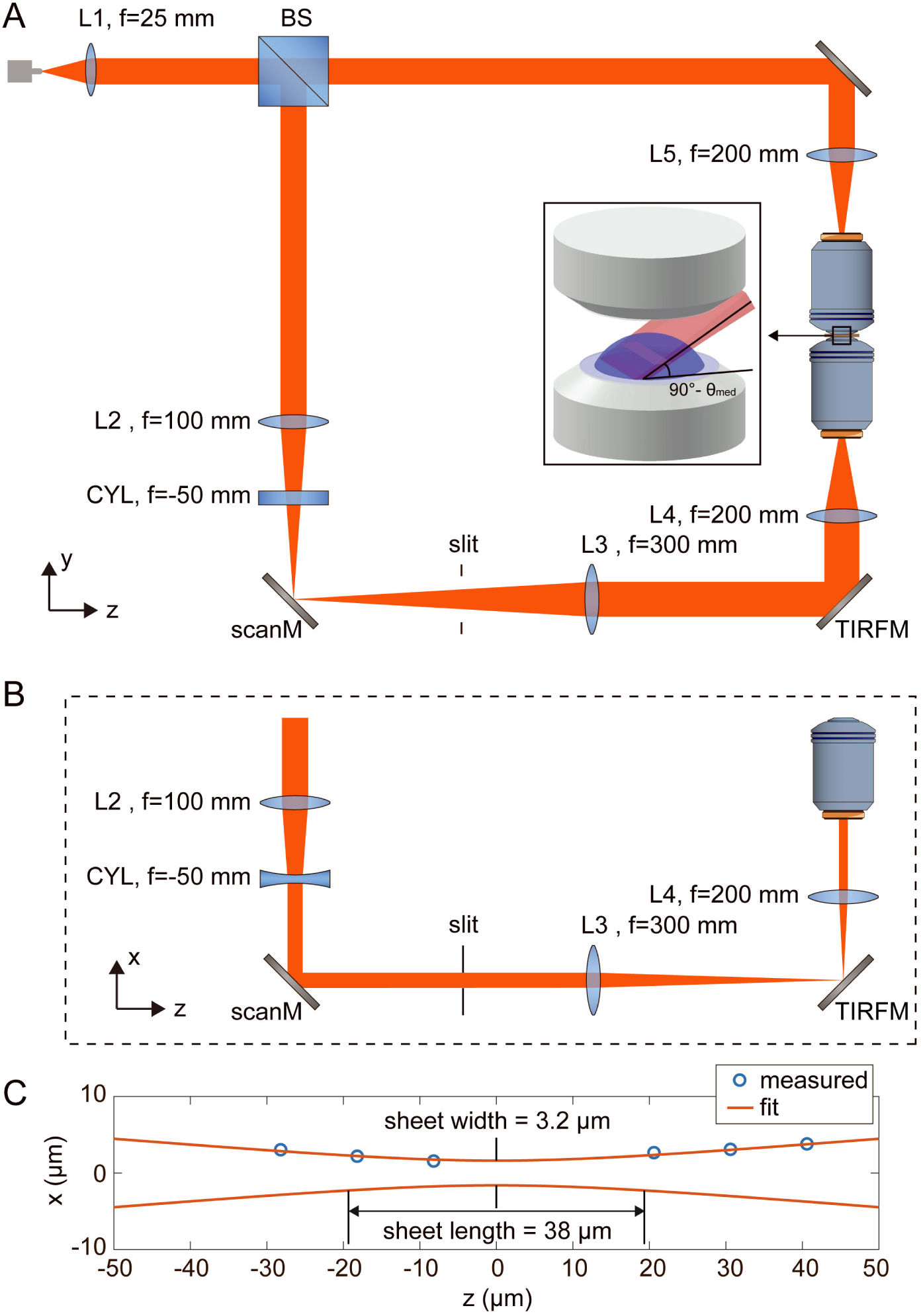
Schematic drawings of Hi-LS (highly-inclined light-sheet) setup and the light-sheet profile. (**A**) Illustration of Hi-LS excitation path in the 4Pi-SMSN system, including an upper epi illumination path and a lower light-sheet illumination path. (**B**) The lower excitation path in x-z plane. (**C**) The light-sheet profile measured at the position that the light-sheet propagation direction was perpendicular to the sample plane. A scan mirror (scanM) at the conjugate pupil plane of the objective was used to translate the light-sheet illumination area. A TIRF mirror (TIRFM) at the conjugate image plane of the sample was used to adjust the light-sheet illumination angle (*θ*_med_) (**note S9**). L: Lens, CYL: cylindrical lens, BS: 50/50 beam splitter, f: focal length.

**Fig S9.**
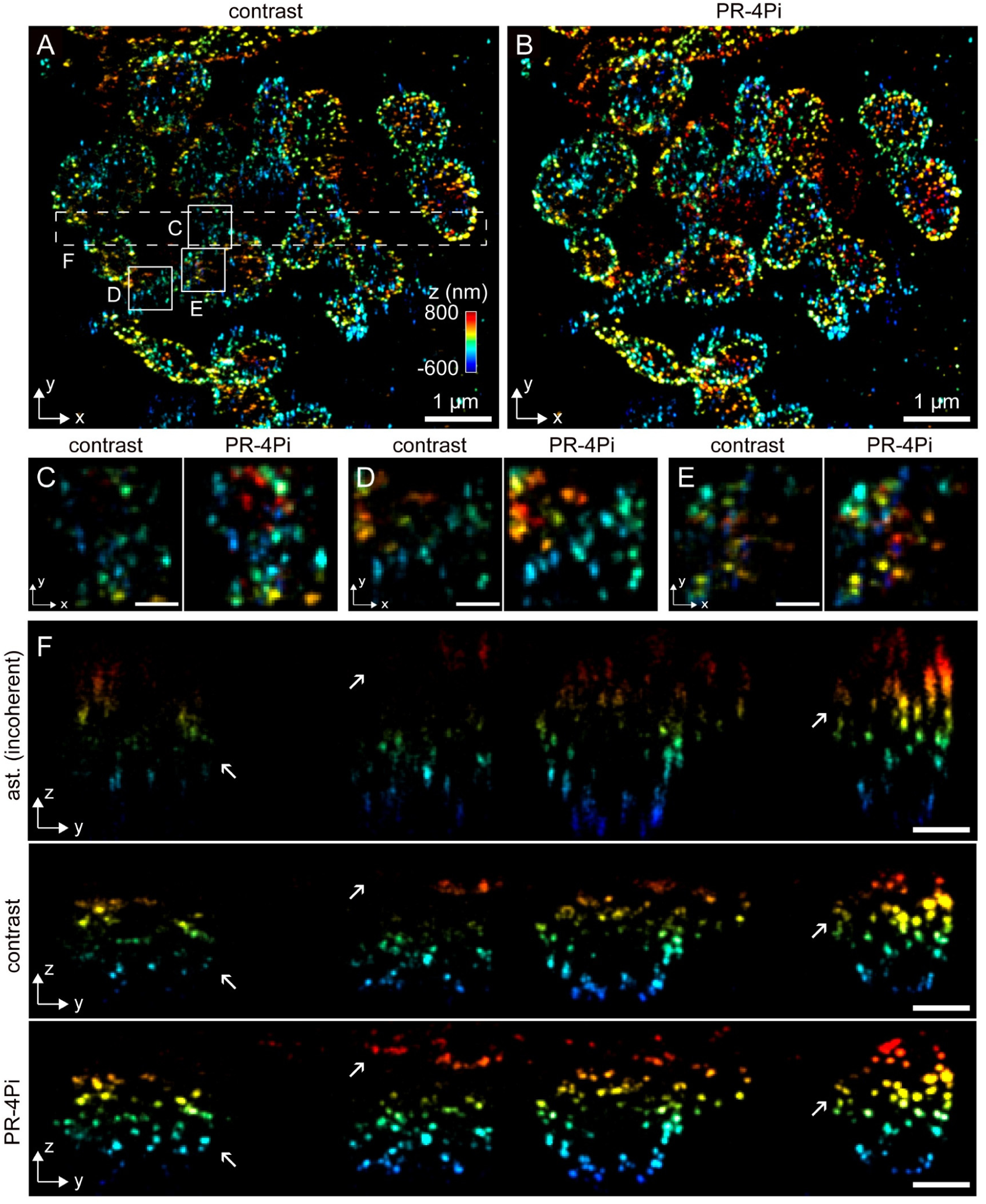
SMSN reconstruction of single-plane imaging of mitochondria (TOM20) in COS-7 cells. (**A, B**) Lateral view of the SMSN reconstruction with PR-4Pi and contrast based methods. (**C-E**) Zoom-in regions in (A, white boxes) at the x-y plane. (**F**) Zoom-in regions in (A, white box with dash lines) at the y-z plane. The astigmatism method (ast.) is based on a Gaussian PSF model and operated on incoherent PSFs that are the sum of the 4Pi-PSFs of the four channels, equivalent to data obtained from the dual-objective 3D imaging method (*12*). Localization results from PR-4Pi show improved resolutions in both lateral and axial dimensions, higher acceptance rate and extended axial localization range. Scale bars: 200 nm in (C-E), 400 nm in (F).

**Fig S10.**
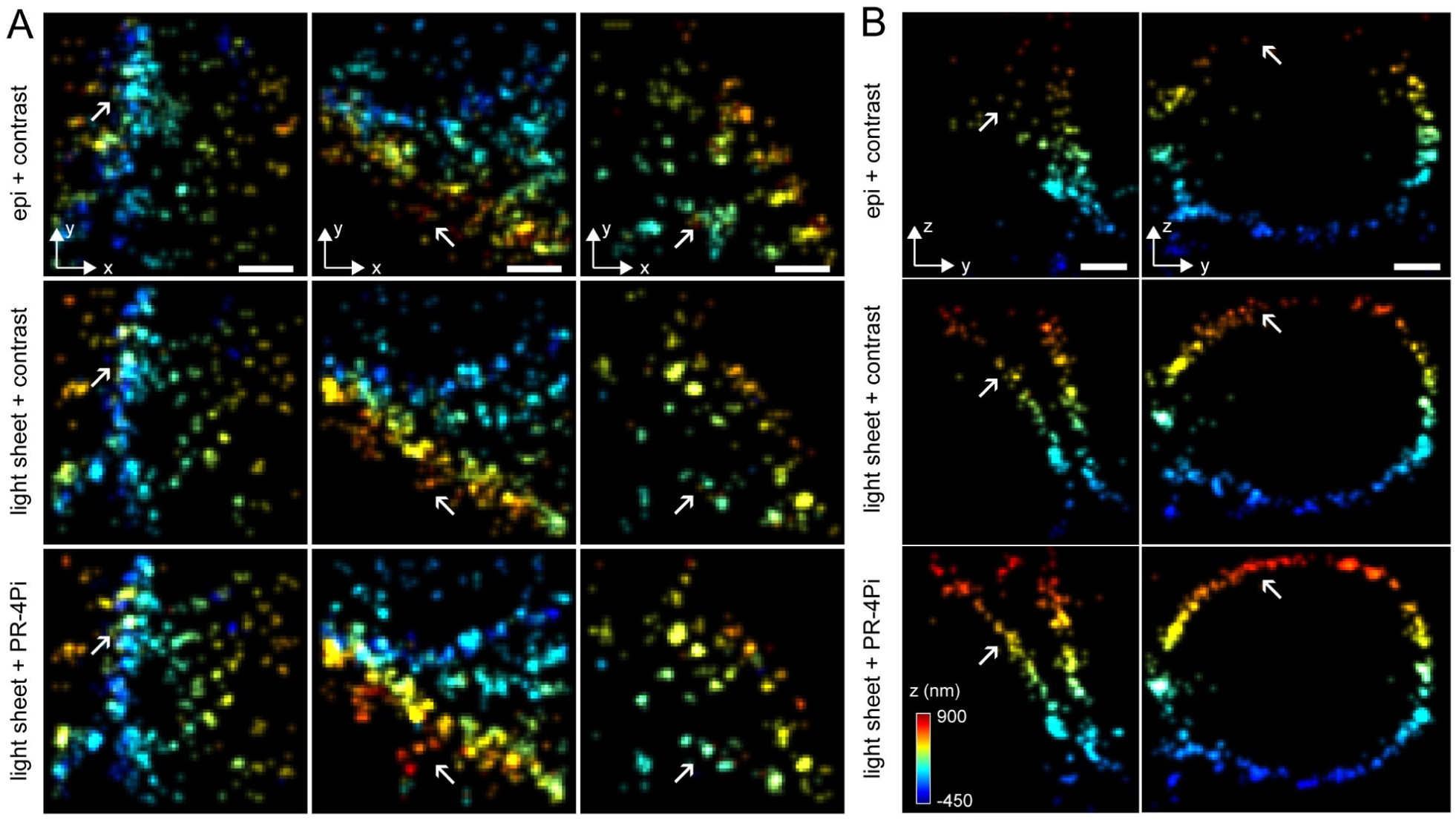
Comparison of SMSN reconstruction of mitochondria (TOM 20) in COS-7 cells with different imaging modalities and analyzing methods. (**A, B**) SMSN reconstructions at the x-y and y-z planes. Data from epi illumination were analyzed with contrast based method, while data from light-sheet illumination were analyzed with both contrast and PR-4Pi method. The sample was imaged at a single axial plane with 20,000 frames under epi illumination followed by 20,000 frames under light-sheet illumination (**Fig. 6** and **fig. S8**). Scale bars: 200 nm.

**Fig S11.**
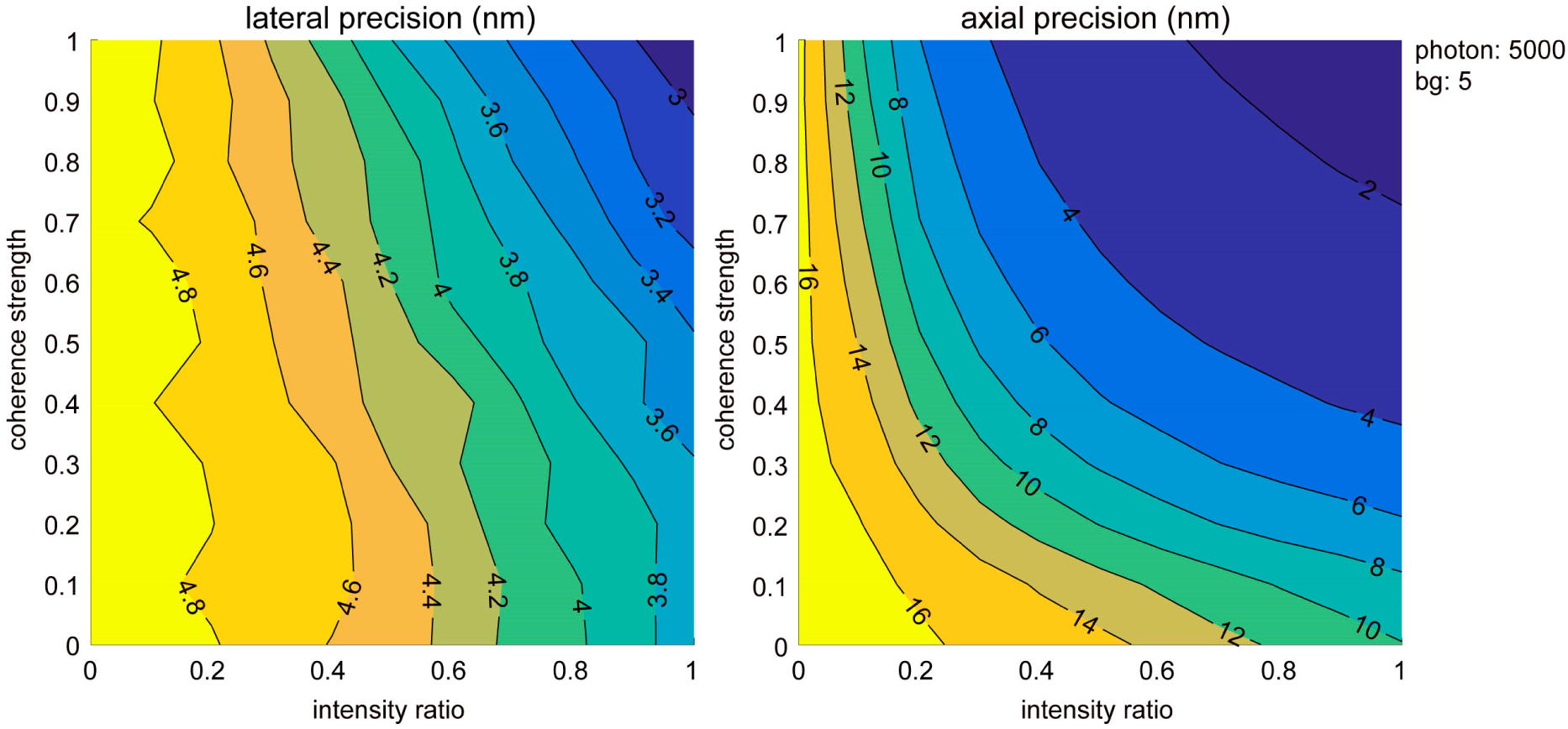
Effect of coherence strength and intensity ratio on lateral and axial localization precisions. Theoretical precisions based on CRLB (**Main Text**) were calculated from 4Pi-PSF models with various coherence strength and intensity ratios (**note S4**). The precision value under each condition was the average of 1000 calculations at *z* positions sampled randomly with a uniform distribution from -0.8 to 0.8 µm.

**Fig S12.**
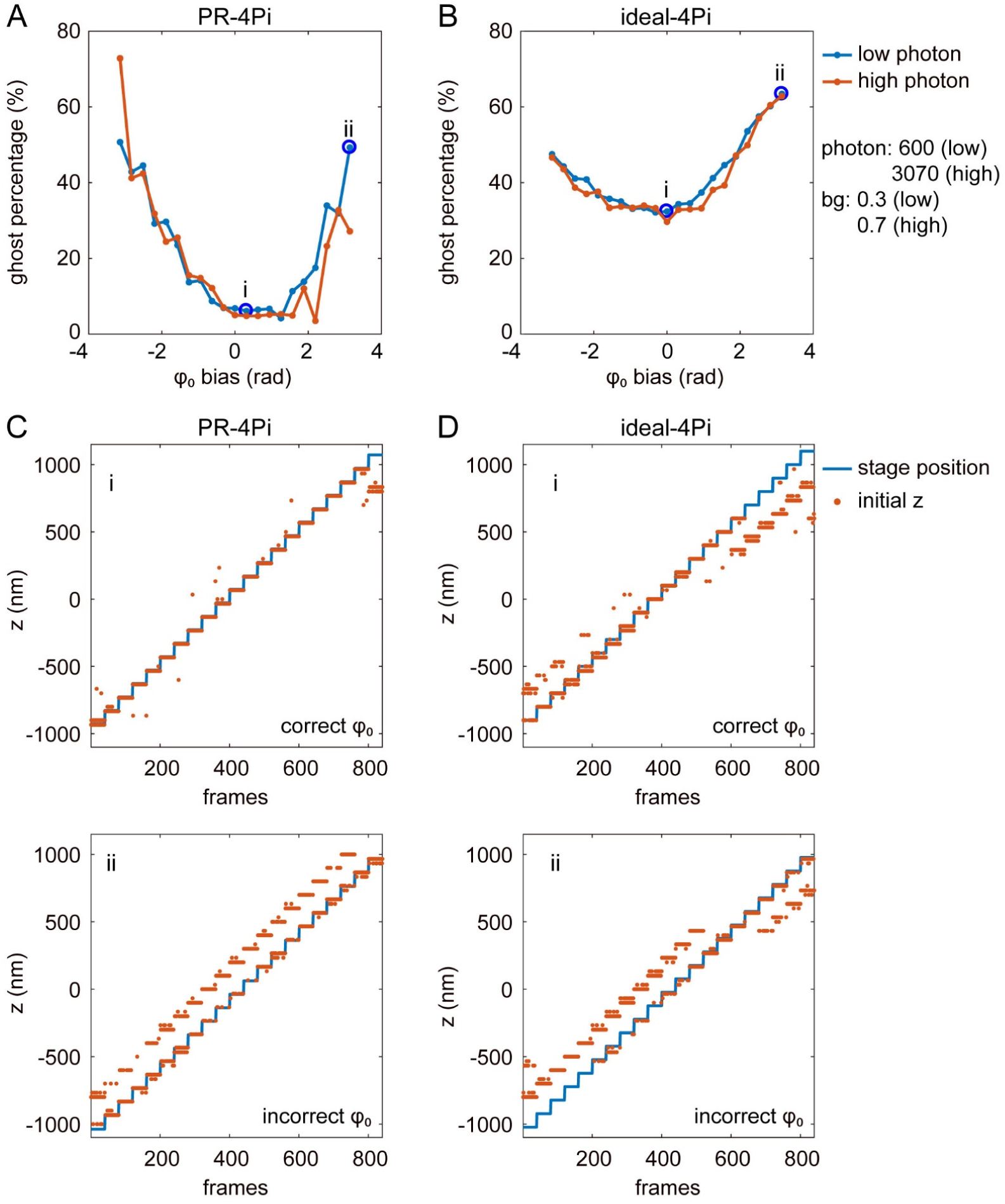
Effect of cavity phase bias on the percentage of ghost localizations. The PR-4PiPSF models were generated from phase retrieved pupil functions and the ideal-4PiPSF models used unaberrated pupil functions and assumed no transmission loss and beam splitting inequality presented in the imaging system. (**A, B**) Ghost percentage from PR and ideal 4Pi-PSF models with different cavity phase (*φ*_0_) biases. (**C, D**) *z* initial guessed positions of the bead data using the PR and ideal 4Pi-PSF models at small and large cavity phase biases (blue circles in A and B). The ghost percentage is defined as the percentage of the initial *z* positions (red dots) at a distance greater than 200 nm from the corresponding stage positions (blue lines). Data were acquired by imaging a fluorescence bead at axial positions from -1 to 1 µm by translating the sample stage with a step size of 100 nm, 40 frames were captured at each axial position (**Materials and Methods**). The estimation results from PR-4Pi method yielded a mean total photon of 600 (3070) per objective and a background (bg) photon of 0.3 (0.7) per pixel for bead data with low (high) signal to noise ratio (SNR).

**Fig S13.**
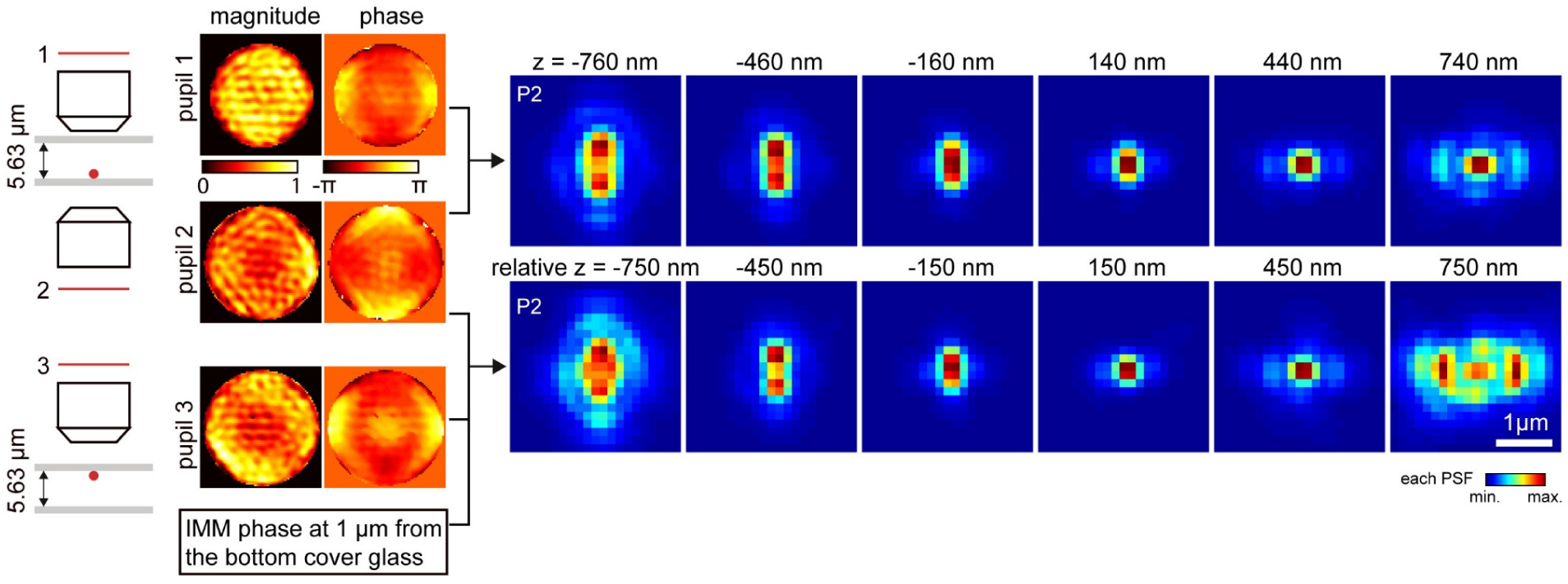
PR-4PiPSF models considering index mismatch aberration (IMM). Two methods are presented here (**note S3**). Method one uses pupil 1 and pupil 2, retrieved independently through the upper and lower objectives by imaging a bead on the bottom cover glass. Pupil 1 includes the index mismatch aberration caused by imaging through the entire sample medium (e.g. 5.63 µm thick). Method two uses pupil 2 and pupil 3, where pupil 3 is retrieved by imaging a bead on the top cover glass through the upper objective. Method 1 is suitable for small imaging depth, less than 4 µm, while method 2 is designed to model emission patterns at arbitrary axial locations inside the sample medium given an imaging depth (e.g. 1 µm) measured from the bottom cover glass and the thickness of the sample medium.

**Fig S14.**
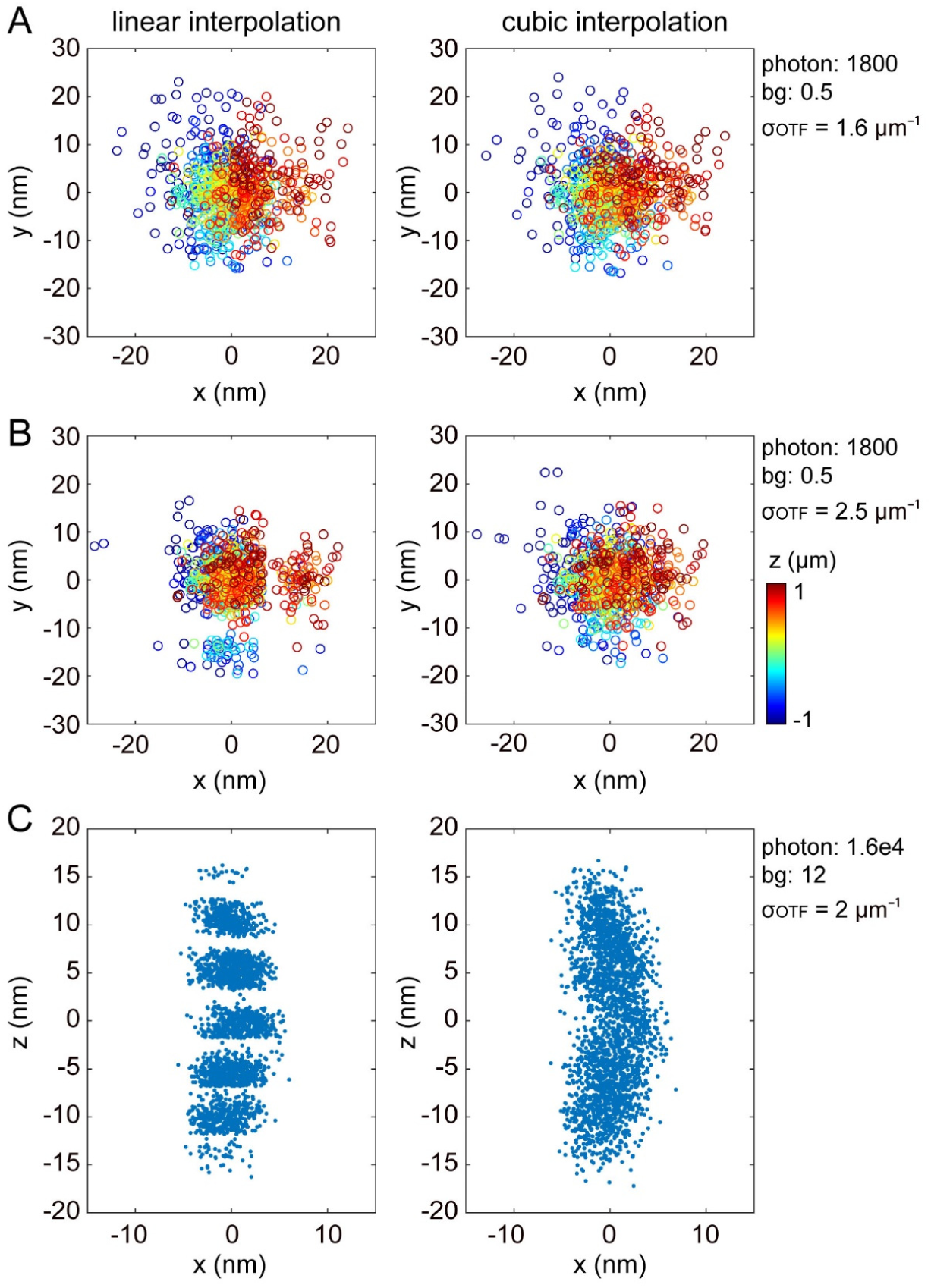
Comparison of linear and cubic interpolation methods on localizations of bead data. (**A, B**) Scatter plots of lateral localizations with linear and cubic interpolated PSF models, smoothed by an OTF rescaling factor (σ_OTF_ equals to *σ*_*kx*_ *= σ*_*ky*_, **note S2**) of 1.6 µm^−1^ and 2.5 µm^−1^ respectively. Individual localization result is color coded according to its axial positions. (**C**) Scatter plots of axial localizations with linear and cubic interpolated PSF models, smoothed by an OTF rescaling factor of 2. We found linear interpolation tends to produce artifacts that are σ_OTF_ dependent in both lateral and axial localizations. Data for testing lateral localizations were acquired by imaging a fluorescence bead at axial positions from -1 to 1 µm by translating the sample stage with a step size of 100 nm, 40 frames were captured at each axial position. Data for testing axial localizations were acquired by imaging an in-focused fluorescence bead for 25 s at 10 Hz. No cavity-phase drift correction was applied for the localization results.

**Fig S15.**
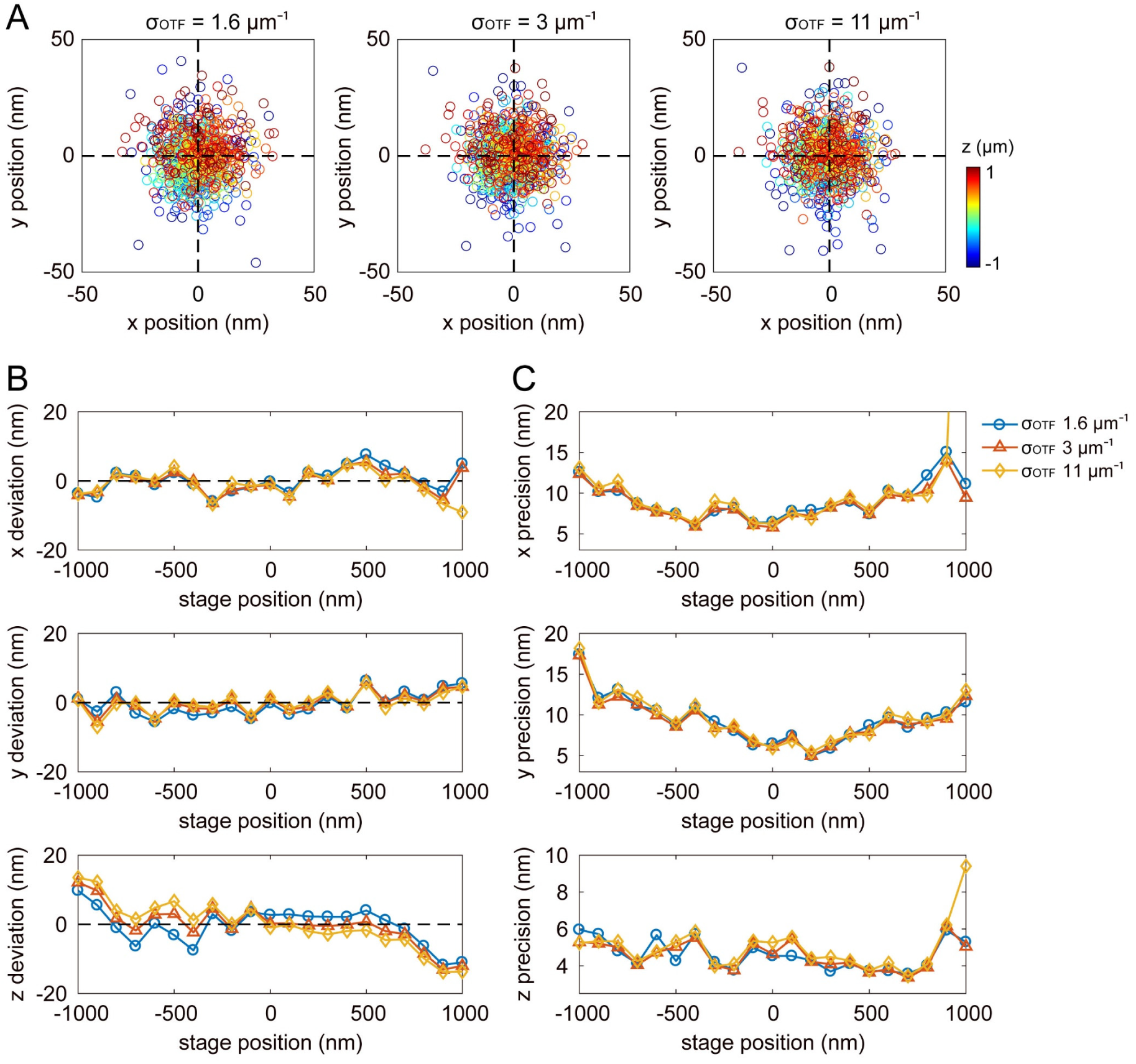
Comparison of localizations of bead data using PSF models smoothed by various OTF rescaling factors. (**A**) Scatter plots of lateral localizations with cubic interpolated PSF models, smoothed by an OTF rescaling factor (*10, 11*) (σ_OTF_ equals to *σ*_*kx*_ *= σ*_*ky*_, **note S2**) of 1.6 µm^−1^, 3 µm^−1^ and 11 µm^−1^ respectively in Fourier space, equivalent to 99.5 nm, 53.0 nm and 14.5 nm in real space. Individual localization result is color coded according to its axial positions. (**B**) Localization deviations in *x, y* and *z*. (**C**) Localization precisions in *x, y* and *z*. Same bead data were used as in **fig. S3**.

**Fig S16.**
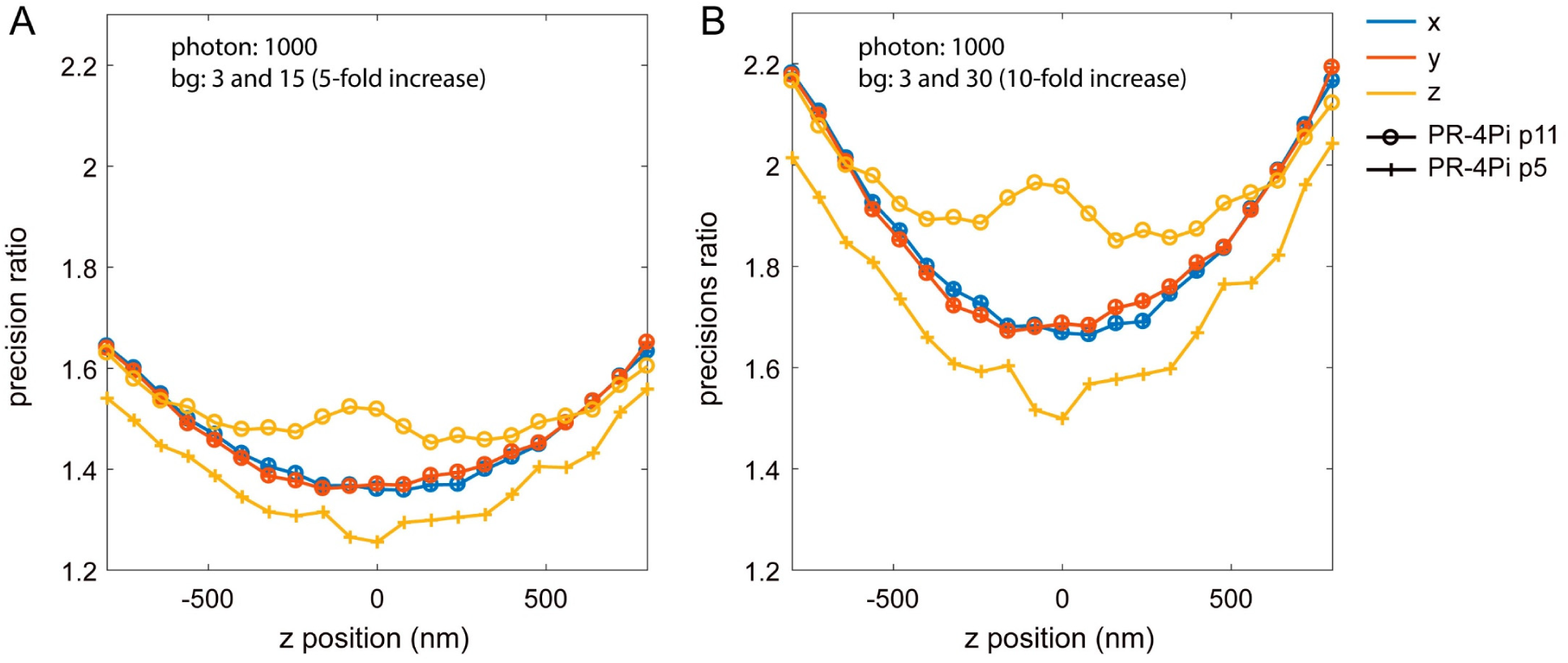
Theoretical improvement of estimation precision with 5 and 10 folds of background reduction. In lateral dimension, PR-4Pi with 5 or 11 parameters improve the precision by the same amount with reduced background. However, for axial localization, PR-4Pi p5 is more susceptible with higher background especially at near focus region, showing less reduced precision. Data were simulated from phase retrieved pupil functions with astigmatism modification and contain 1000 PSFs at each axial position in the range of -900 nm to 900 nm, with a step size of 100 nm.

**Fig S17.**
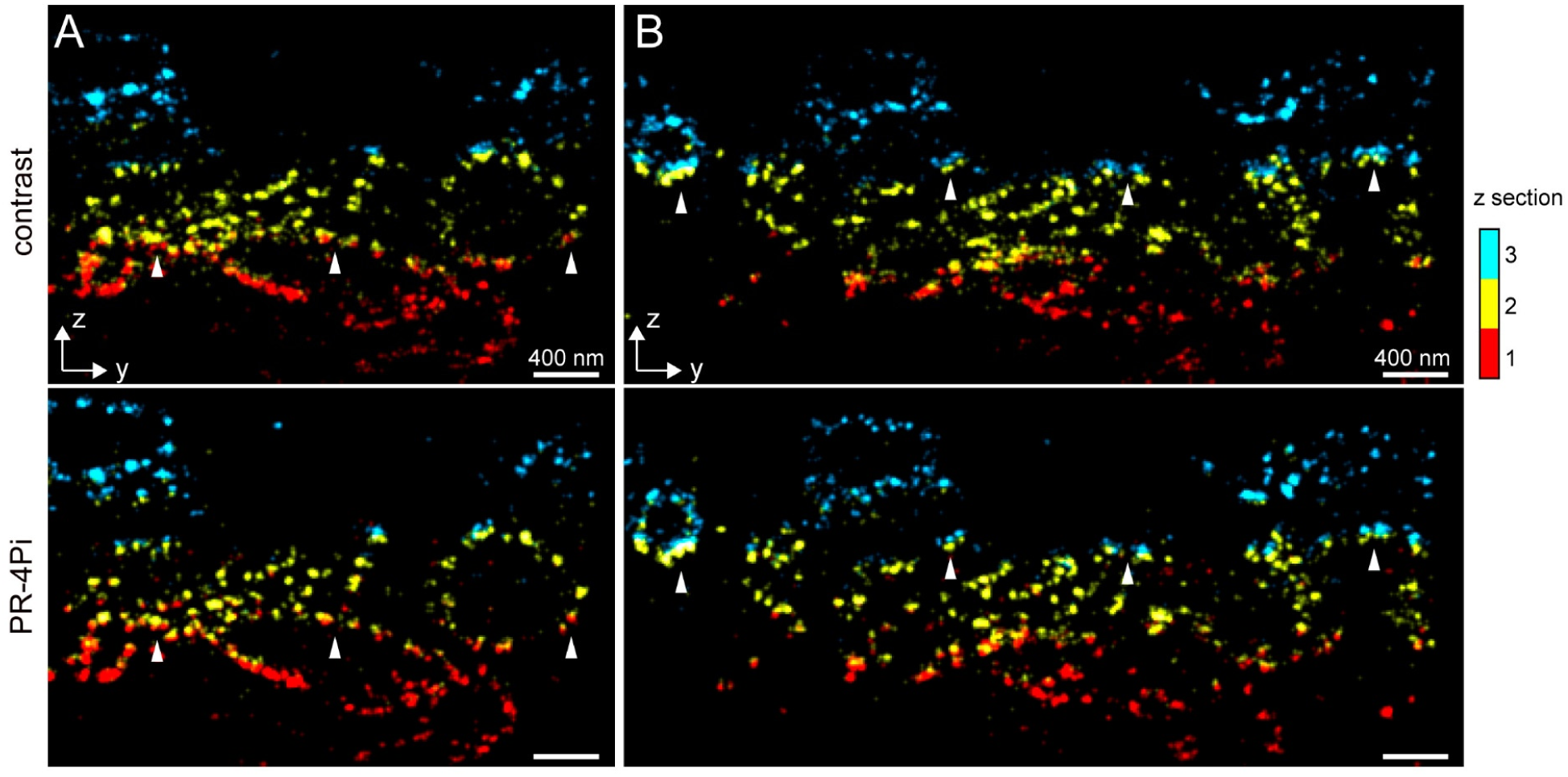
Comparison of section alignment for multi-section imaging of mitochondria in COS-7 cell. (**A**,**B**) SMSN reconstructions at the y-z planes. Color represents optical sections, the step size between consecutive sections is 800 nm. PR-4Pi achieves more accurate alignment between optical sections for its higher localization precision/accuracy and acceptance rate as well as extended axial range of each optical section.

**Fig S18.**
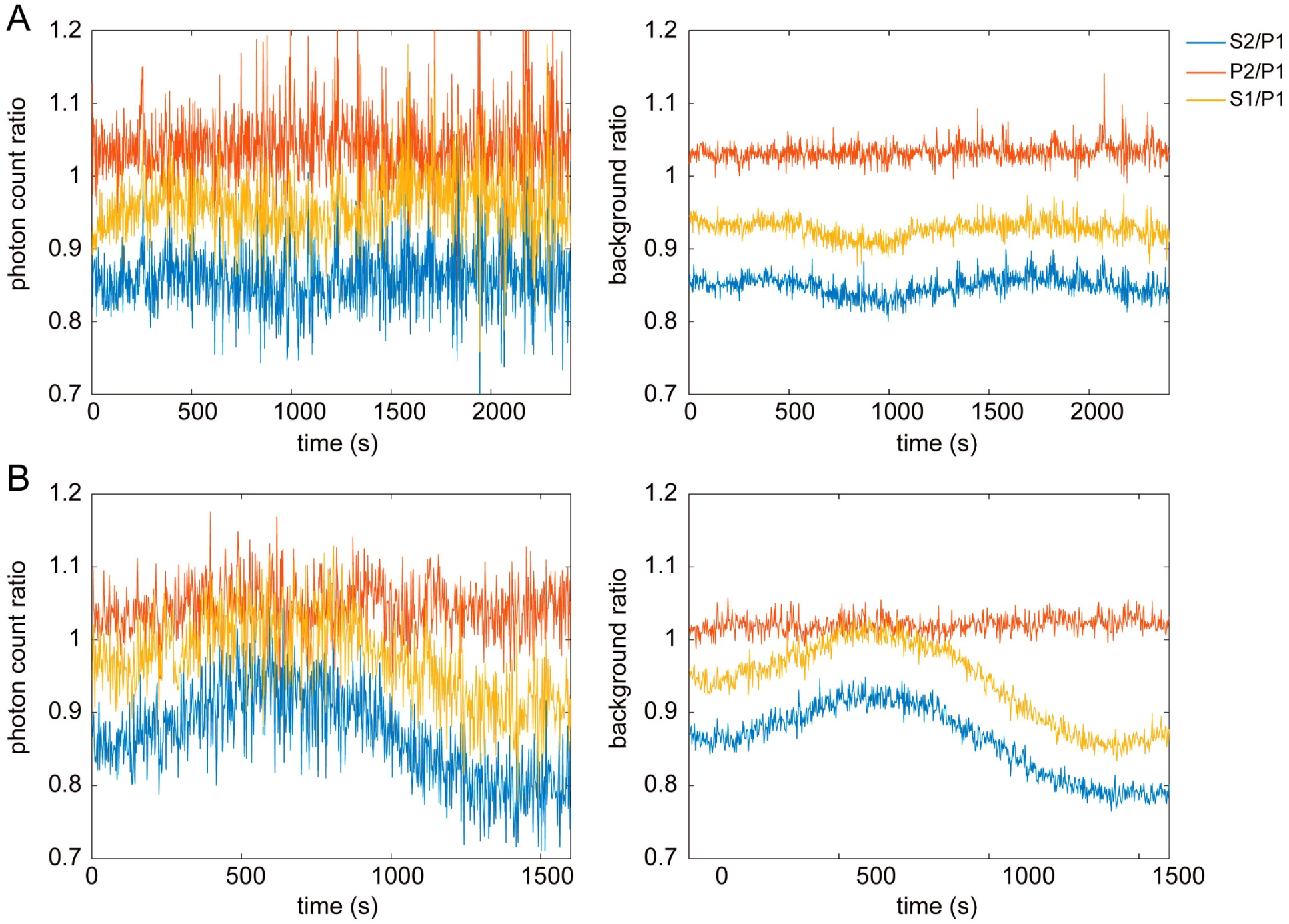
Estimated photon and background ratios between four detection channels from mitochondria imaging in COS-7 cell. (**A**) Photon/background ratios of multi-section imaging results from **Fig. 5**. (**B**) Photon/background ratios of single-section imaging results from **fig. S9**. Ratios were calculated from the averaged photon/background values of every 100 frame from four detection channels: P1, S2, P2 and S1 (**Fig. 1**). The ratio traces show a fluctuation of intensity ratio between s-and p-polarizations over the acquisition time. We suspect this fluctuation might be caused by the changes of conditions of the imaging buffer environment or the cell samples. Therefore, to account for the sample specific change of intensity ratio between s-and p-polarizations, we proceeded using the PR-4Pi localization algorithm with 11 estimation parameters to ensure accurate (smaller bias) localization results.

**Table S1.**
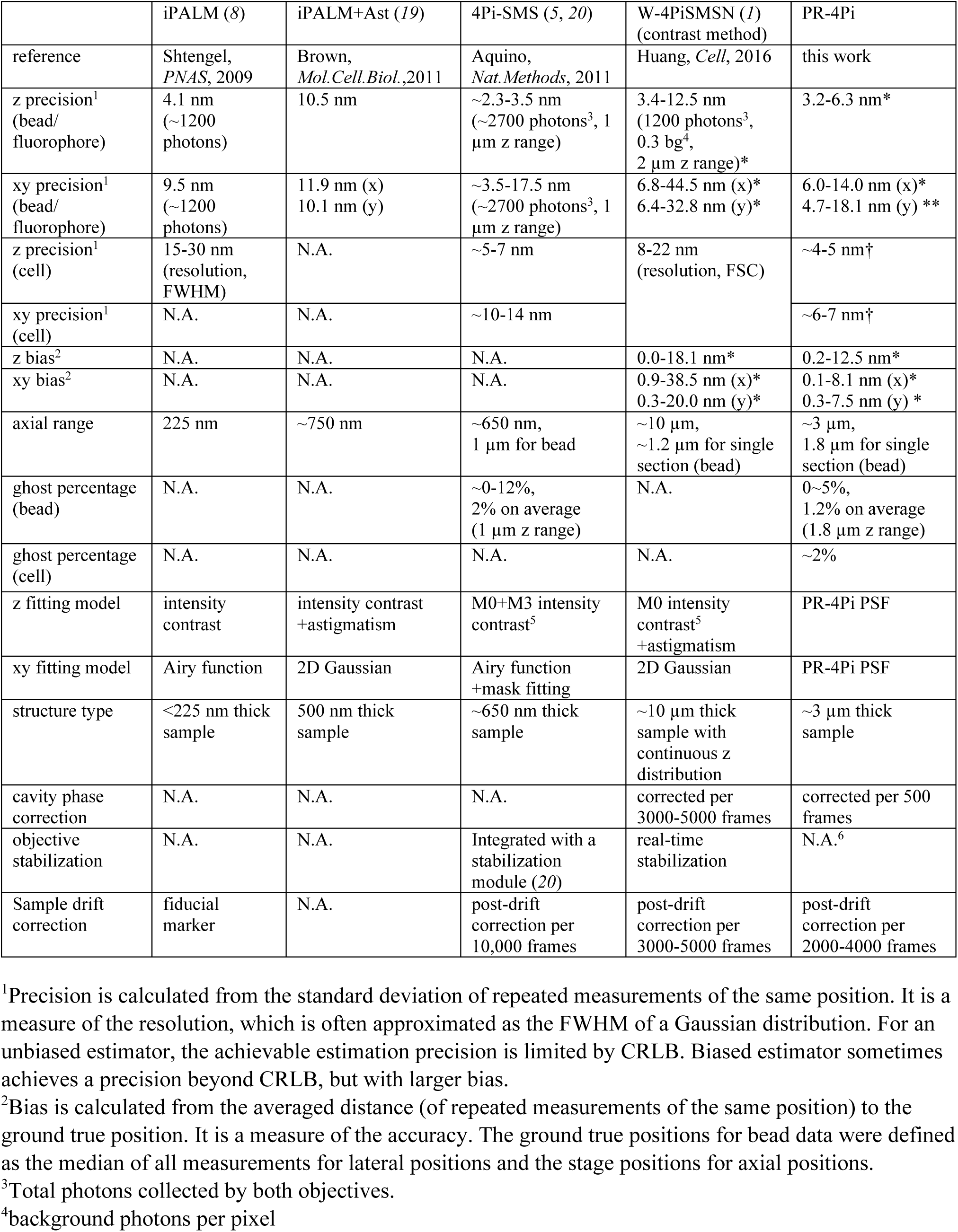
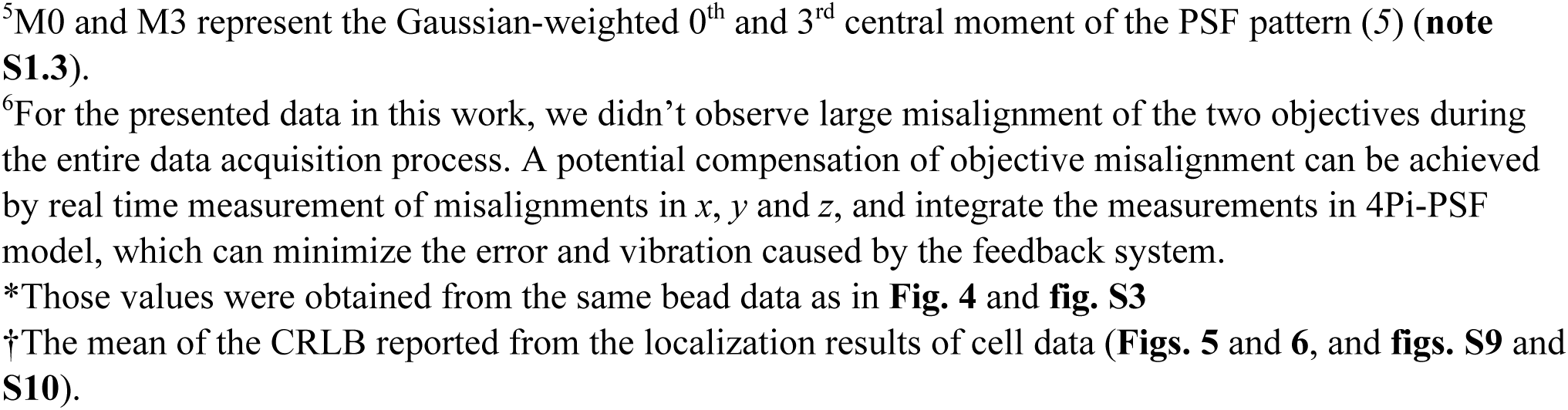
Comparison of interferometric single-molecule localization microscopies

|  | iPALM (8) | iPALM+Ast (19) | 4Pi-SMS (5, 20) | W-4PiSMSN (1)<br>(contrast method) | PR-4Pi |
| --- | --- | --- | --- | --- | --- |
| reference | Shtengel,<br>*PNAS*, 2009 | Brown,<br>*Mol. Cell. Biol.*, 2011 | Aquino,<br>*Nat. Methods*, 2011 | Huang, *Cell*, 2016 | this work |
| z precision<sup>1</sup><br>(bead/<br>fluorophore) | 4.1 nm<br>(~1200<br>photons) | 10.5 nm | ~2.3-3.5 nm<br>(~2700 photons<sup>3</sup>, 1<br>μm z range) | 3.4-12.5 nm<br>(1200 photons<sup>3</sup>,<br>0.3 bg<sup>4</sup>,<br>2 μm z range)* | 3.2-6.3 nm* |
| xy precision<sup>1</sup><br>(bead/<br>fluorophore) | 9.5 nm<br>(~1200<br>photons) | 11.9 nm (x)<br>10.1 nm (y) | ~3.5-17.5 nm<br>(~2700 photons<sup>3</sup>, 1<br>μm z range) | 6.8-44.5 nm (x)*<br>6.4-32.8 nm (y)* | 6.0-14.0 nm (x)*<br>4.7-18.1 nm (y) ** |
| z precision<sup>1</sup><br>(cell) | 15-30 nm<br>(resolution,<br>FWHM) | N.A. | ~5-7 nm | 8-22 nm<br>(resolution, FSC) | ~4-5 nm† |
| xy precision<sup>1</sup><br>(cell) | N.A. | N.A. | ~10-14 nm |  | ~6-7 nm† |
| z bias<sup>2</sup> | N.A. | N.A. | N.A. | 0.0-18.1 nm* | 0.2-12.5 nm* |
| xy bias<sup>2</sup> | N.A. | N.A. | N.A. | 0.9-38.5 nm (x)*<br>0.3-20.0 nm (y)* | 0.1-8.1 nm (x)*<br>0.3-7.5 nm (y) * |
| axial range | 225 nm | ~750 nm | ~650 nm,<br>1 μm for bead | ~10 μm,<br>~1.2 μm for single<br>section (bead) | ~3 μm,<br>1.8 μm for single<br>section (bead) |
| ghost percentage<br>(bead) | N.A. | N.A. | ~0-12%,<br>2% on average<br>(1 μm z range) | N.A. | 0~5%,<br>1.2% on average<br>(1.8 μm z range) |
| ghost percentage<br>(cell) | N.A. | N.A. | N.A. | N.A. | ~2% |
| z fitting model | intensity<br>contrast | intensity contrast<br>+astigmatism | M0+M3 intensity<br>contrast<sup>5</sup> | M0 intensity<br>contrast<sup>5</sup><br>+astigmatism | PR-4Pi PSF |
| xy fitting model | Airy function | 2D Gaussian | Airy function<br>+mask fitting | 2D Gaussian | PR-4Pi PSF |
| structure type | <225 nm thick<br>sample | 500 nm thick<br>sample | ~650 nm thick<br>sample | ~10 μm thick<br>sample with<br>continuous z<br>distribution | ~3 μm thick<br>sample |
| cavity phase<br>correction | N.A. | N.A. | N.A. | corrected per<br>3000-5000 frames | corrected per 500<br>frames |
| objective<br>stabilization | N.A. | N.A. | Integrated with a<br>stabilization<br>module (20) | real-time<br>stabilization | N.A.<sup>6</sup> |
| Sample drift<br>correction | fiducial<br>marker | N.A. | post-drift<br>correction per<br>10,000 frames | post-drift<br>correction per<br>3000-5000 frames | post-drift<br>correction per<br>2000-4000 frames |

<sup>1</sup>Precision is calculated from the standard deviation of repeated measurements of the same position. It is a measure of the resolution, which is often approximated as the FWHM of a Gaussian distribution. For an unbiased estimator, the achievable estimation precision is limited by CRLB. Biased estimator sometimes achieves a precision beyond CRLB, but with larger bias.
<sup>2</sup>Bias is calculated from the averaged distance (of repeated measurements of the same position) to the ground true position. It is a measure of the accuracy. The ground true positions for bead data were defined as the median of all measurements for lateral positions and the stage positions for axial positions.
<sup>3</sup>Total photons collected by both objectives.
<sup>4</sup>background photons per pixel
<sup>5</sup>M0 and M3 represent the Gaussian-weighted 0<sup>th</sup> and 3<sup>rd</sup> central moment of the PSF pattern (5) (**note S1.3**).
<sup>6</sup>For the presented data in this work, we didn't observe large misalignment of the two objectives during the entire data acquisition process. A potential compensation of objective misalignment can be achieved by real time measurement of misalignments in $x$ , $y$ and $z$ , and integrate the measurements in 4Pi-PSF model, which can minimize the error and vibration caused by the feedback system.
\*Those values were obtained from the same bead data as in **Fig. 4** and **fig. S3**
†The mean of the CRLB reported from the localization results of cell data (**Figs. 5** and **6**, and **figs. S9** and **S10**).

